# Neonatal liver niches program T cell tolerance

**DOI:** 10.64898/2026.01.13.698576

**Authors:** Eva-Lena Stange, Trong-Hieu Nguyen, Dustyn Mendoza, Aiara Lobo Gomes, Marlene Sophia Kohlhepp, Jonas Pes, Shahed Al Bounny, Julian Brueck, Yunus Cetiner, Urs Moerbe, Milas Ugur, Cristina Kalbermatter, Jarrett Lopez-Scarim, Aline Dupont, Susan A. V. Jennings, Julia Heckmann, Aaron Silva-Sanchez, Solveig Runge, Christoph Kuppe, Oliver Pabst, Thomas Clavel, Dorothee Viemann, Stephan P. Rosshart, Vuk Cerovic, Stephanie C. Ganal-Vonarburg, Adrien Guillot, Eva Billerbeck, Mathias W. Hornef, Natalia Torow

## Abstract

After birth, the immune system must learn to tolerate a rapidly changing milieu of commensals and self while remaining ready for pathogens. Here we characterize the neonatal liver as a central hub in this process: In postnatal week 1–2, the liver hosts a developmentally encoded, microbiota-independent expansion of regulatory T cells (Tregs) that coexists with microbiota-tuned conventional wave of activated CD4⁺ T cells (Tconvs). Mechanistically, the Treg expansion is governed by MHCII-mediated antigen presentation by CCR7+ cDC1s, which establish tolerogenic DC:T cell clusters in the liver parenchyma, allowing for local expansion and control via PD-L1 checkpoints that selectively increase Tregs without unleashing Tconvs. Importantly, this transient, neonatal program predisposes hepatotropic viral infections to progress toward chronic disease but also protects the adult liver from steatotic disease. These data position the neonatal liver as a unique site of early life T-cell education with timing-sensitive implications for early-life interventions.

## Introduction

Birth marks a distinct phase for adaptive immunity in the murine host: peripheral tissues are rapidly seeded by thymus-derived recent thymic emigrants (RTEs), and the early peripheral T cell pool is strongly enriched for cells that are more self-reactive and less stringently tuned than adult naïve T cells (*1–4*). At the same time, microbial colonization takes hold at barrier surfaces (*5, 6*). Therefore, the neonatal immune system must enforce peripheral self-tolerance and, in parallel, establish tolerance to newly encountered commensals.

Consequently, early life immunity is characterized by regulatory T cell (Treg) waves, although the timing and mechanism of induction differ by tissue. In the intestine, T cells remain naïve and have restricted tissue access until weaning, when a major dietary transition arrives with solid food, and an intestine-specific, microbiota- and dietary antigen-dependent Treg burst has been described (*7–9*). This gut-specific Treg burst is induced by the maturation and development of intestinal antigen uptake mechanisms at weaning, including goblet cell–associated passages and M-cell–mediated transcytosis (*10–13*). Although early-life Treg expansion has been most intensively studied in the gut, pre-weaning Treg waves have also been documented in lung and skin, where transient, microbiota-dependent expansions curb inflammatory priming (*14–16*). Together, these observations raise a broader question: where and how is tolerance enforced prior to the maturation of classical intestinal sampling?

In the liver, a pre-weaning Treg wave has been noted previously (*17, 18*), implicating the liver as a key tolerogenic environment in early life. Further, in the adult liver, antigen presentation and coinhibitory pathways bias adaptive immunity towards deletion, anergy, or Treg induction (*19–23*). Moreover, the liver can also organize structured immune microenvironments in inflammatory settings, underscoring its capacity to support local immune organization (*19, 21, 24*). However, the mechanisms of induction, antigen presentation and microbiota dependence as well as the functional relevance of the early-life hepatic Treg wave remain largely unclear. Elucidating these tolerogenic molecular pathways represents an urgent need for the rational design of effective vaccines and therapies targeting neonatal infections.

Here, we define an early-life immune architecture in which the neonatal liver functions as a transient, tolerogenic CD4⁺ T cell niche. We demonstrate that the neonatal hepatic Treg wave represents a developmentally programmed, microbiota-independent, locally controlled Treg expansion that occurs concurrently with a microbiota-mediated Tconv activation. Crucially, we identify CCR7⁺ type 1 conventional dendritic cells (cDC1s) as the dominant antigen presenting cell (APC) population that anchors Treg- and Tconv-containing clusters within the neonatal liver parenchyma. Furthermore, we show the key role of cDC1-associated PD-L1 signaling in shaping the balance between Treg accumulation and Tconv activation, providing a mechanism by which the neonatal liver calibrates responses to portal-derived microbial constituents. Finally, we link this early-life tolerogenic set-point to delayed control of pediatric hepatotropic virus infection and increased resistance to metabolic-induced liver inflammation in the adult host.

### A developmentally-programmed postnatal hepatic Treg wave constrains microbiota-dependent Tconv expansion and activation

A comparison of the CD4^+^ T cell compartment in early life and adulthood by flow cytometry revealed a notable enrichment of Tregs in the neonatal liver of specific-pathogen-free (SPF) mice, in marked contrast to other peripheral or lymphoid tissues (Fig. 1a) and in accordance with the previously reported “neonatal Treg wave” (*17, 25*). Likewise, the total hepatic CD4 and CD8 T cell density was higher in the neonatal liver but not spleen or small intestinal *lamina propria* (SILP) (Figs. S1A-B). To further explore the inductive mechanisms and functional implications of this early T cell/Treg wave, we performed a detailed kinetic analysis across the pre-weaning period (Extended data 1c-e). The hepatic CD4⁺ T cell compartment underwent a pronounced early-life expansion that peaked at postnatal day (PND) 7-9 (Fig. S1c). Within this wave, Tregs peaked at PND7, accounting for up to ∼35 % of hepatic CD4⁺ T cells (Fig. 1 b; Extended data 1d). Conventional CD4⁺ T cells (Tconv) expanded with slightly delayed kinetics, peaking at PND9 before both populations gradually declined toward adult set points (Fig. 1b). Liver CD8^+^ T cells showed a similar kinetic to CD4^+^ T cells, while non-classical T cell populations, such as TCRγδ^+^ T cells and NKT cells also showed substantial expansion in the pre-weaning period, albeit with distinct kinetics (Fig. S1e). Immunofluorescence imaging of hepatic tissue confirmed the increased density of T cells along with Tregs in neonatal livers and revealed clusters of co-localizing Tregs with other T cell subsets distributed throughout the liver parenchyma including periportal areas (Fig. S1c-d). In adult livers, T cell clusters were present at almost exclusively confined to periportal areas (Fig. S1f) and were less diverse in composition (Fig. 1c). Collectively, these data point to the neonatal liver as a key site of T cell immune activation and regulation in early life.

**Figure 1:**
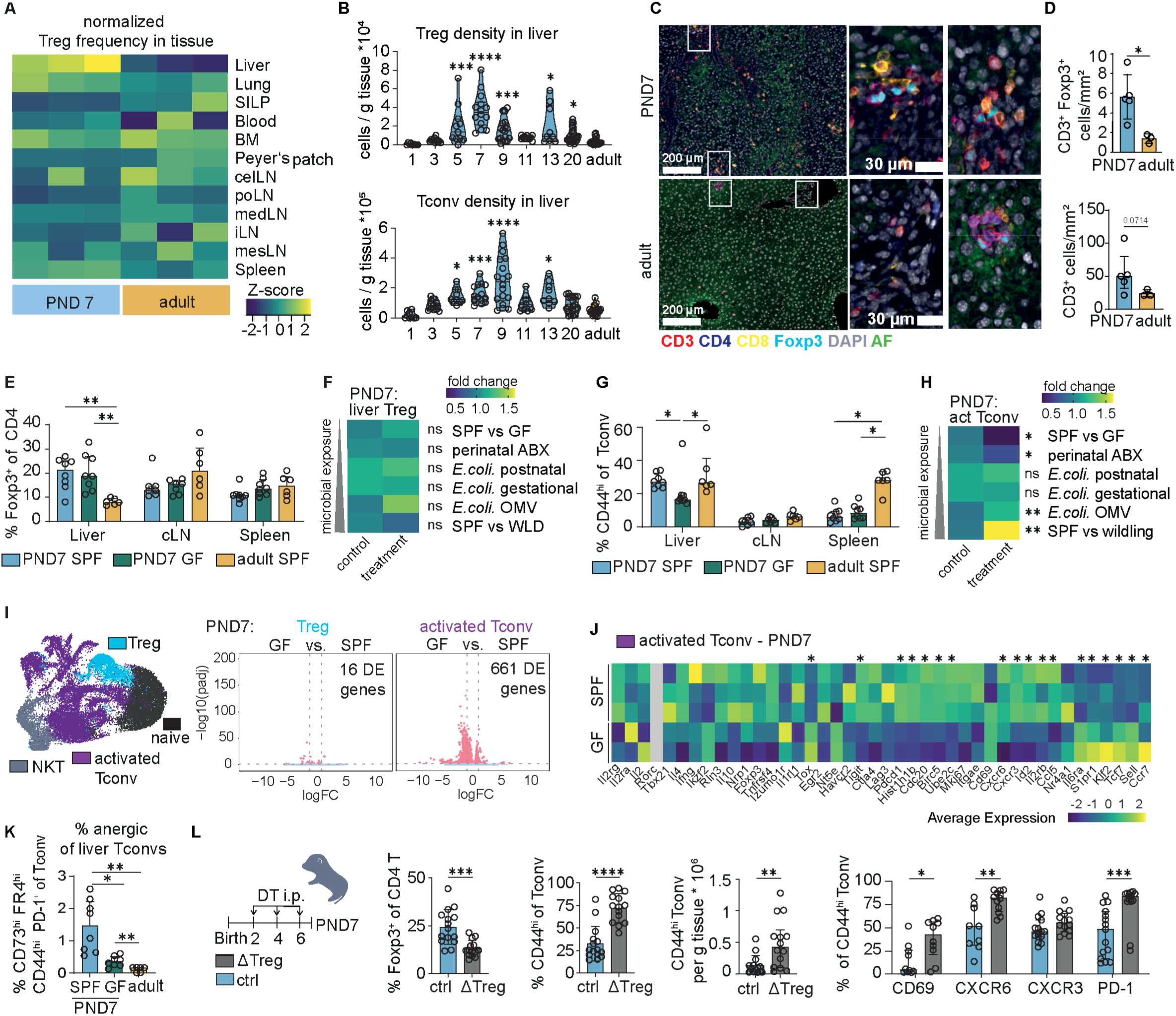
A transient Treg peak restrains microbiota-tuned Tconv activation across hepatic clusters. (A) Row z-score normalized data % of Treg within the CD4 T cell compartment in liver, lung, small intestinal lamina propria (siLP), blood, bone marrow (BM), Peyer’s Patch, celiac lymph node (cLN), portal lymph node (pLN), mediastinal lymph node (medLN), inguinal lymph node (iLN) and mesenteric lymph node (mLN) and spleen in adult and 7-day old neonatal mice. Flow cytometry, 3 mice per group, Representative data from 1-5 experiments. (B) Foxp3^+^ CD4 T cells (Treg) and Foxp3^-^ CD4T cells (Tconv) per g liver tissue over the pre-weaning period. Violin plots of 6-20 mice per timepoint from 2-6 independent experiments. Kruskal Wallis test + post hoc Dunn’s comparison of each lime point to the adult group. (C) lmmunofluorescence of T cell markers (CD3, CD4, CD8*α*, Foxp3) in neonatal (PND7) and adult liver tissue. (D) Automated full-slide quantification of in (c): CD3^+^ and Foxp3^+^ neonatal and adult liver tissue from 3-5 mice group. Welch I-tests. (E) % of Treg within CD4 T cell compartment in liver, celiac lymph node (cLN) and spleen (Spl) of neonatal (PND7) SPF and GF mice and adult SPF ice. 6-8 mice per group from two independent pooled experiments. Brown-Forsythe ANOVA + Dunnetrs multiple comparison test (liver, cLN) or One Way ANOVA + post hoc Holm-Sidak’s multiple test (Spl). (F) Heatmap showing the mean fold change of % Treg of liver CD4 T cells normalised to the median of the respective control group (untreated/mock-treated SPF mice for comparison to GF, AVNM, OMV and WLD; untreated GF mice for GC and PC) of the different microbial colonization experiments shown in Ext. Data 2a. Asterisks represent the results of the statistical tests of the % of Treg data from Ext.data 2a-e . (G) % of CD44^hi^ Tconv within CD4 T cell compartment in liver, celiac lymph node (cLN) and spleen (Spl) of neonatal (PND7) SPF and GF mice and adult SPF mice. 6-8 mice per group from two independent pooled experiments. Kruskal Wallis + Dunn’s multiple comparison test (Liver, spleen) or One Way ANOVA + post hoc Holm-Sidak’s multiple test (cLN). (H) Heatmap showing the mean fold change of % CD44^hi^ liver Tconv in the liver normalized to the median of the respective control group (untreated/mock-treated SPF mice for comparison to GF, AVNM, OMV and WLD; untreated GF mice for GC and PC) of the different microbial colonisation experiments shown in Ext. Data 2a. Asterisks represent the results of the statistical tests of the %CD44^hi^ of Tconv data from Ext.data 2a-e. (I) UMAP of liver CD4 T cells from different neonatal and adult lime-points of SPF and GF and volcano plots showing the significant differentially expressed DE genes between 7-day old SPF and GF mice. (J) Expression of a curated list of genes between activated liver Tconvs (purple cluster) in neonatal (PND7) SPF and germ-free mice normalized column expression. Asterisks indicates DE gene between PND7 SPF and PND7 GF mice. (K) Frequency of anergic cells (defined as CD73^hi^ FR4^hi^ PD-1^+^ CD44^hi^ Tconvs) within Tconvs in 7-day-old SPF and GF liver as well as adult SPF liver measured by flow cytometry. 8 mice from 2 experiments. Brown-Forsythe ANOVA + Dunnetrs multiple comparison test. (L) Experimental layout of neonatal Treg depletion experiment; frequency of Treg and CD44^hi^ Tconv in liver CD4 T cells and hepatic density of activated Tconvs measured by CD44 expression and expression of CD69, CXCR6, CXCR3 and PD-1 in CD44^hi^ Tconv. 13-15 mice pooled from 4 independent experiments: Multiple Mann Whitney U tests, Welch I-test ( % Treg of CD4 T cells, % CD44^hi^ of Tconvs) or Mann Whitney U test (CD44^hi^ Tconv density). All experiments containing flow cytometry were done with the compared groups processed on the same day within 1 experiment. Asterisks indicate significance levels: * < 0.05, ** < 0.01, *** < 0.001 **** < 0.0001.

Previous reports have suggested that the neonatal Treg expansion in the liver is dependent on the intestinal microbiota(*17, 25*). To test this, we compared the hepatic T cell compartment of neonatal mice reared under SPF conditions to those of germ-free (GF) neonates. Surprisingly, the hepatic (and liver-draining lymph node) Treg frequency at PND7 remained unchanged (Fig. 1e). To further understand the dependence of neonatal T cells on stimuli from the microbiota and their timing, we analysed litters born after perinatal treatment of dams with broad-spectrum antibiotics, GF mice following “reversible colonization” with auxotrophic *E. coli* HA107 either during pregnancy or in the first week after birth, or neonatal SPF mice treated with *E. coli* outer membrane vesicles (OMVs) for 8h. Additionally, we compared neonatal SPF mice with “wildlings”, which harbour a more diverse microbiota. Strikingly, despite the highly variable strength and timing of microbial exposure, we observed no significant difference in the overall proportion of hepatic Tregs on PND7, demonstrating that the hepatic neonatal Treg wave develops independently of the host microbiota (Fig. 1f; Fig. S2a). In marked contrast, the proportion of neonatal activated CD44^hi^ Tconvs was significantly reduced in the liver but not the spleen or celiac lymph node of GF mice (Fig. 1g) as well as in livers of neonates from antibiotic-treated dams. Conversely, hepatic CD44^hi^ Tconvs showed a significant increase shortly after exposure of SPF neonates to bacterial OMVs as well as in wildlings (Fig. 1h; Figs. S2a-e). By adulthood, activated Tconv frequencies had equalized in germ-free mice, indicating that this distinct neonatal sensitivity to microbial exposure dissipates once the hepatic immune landscape matures (Fig. S2f). Collectively, these data show that, while the hepatic Treg wave is developmentally controlled, the Tconv compartment in the neonatal liver is sensitive to microbial cues.

In order to understand the identified changes in the liver immune landscape at this key period, we performed scRNAseq analysis of hepatic CD45^+^ leukocytes from neonatal and adult SPF and GF mice (Fig. S3). Transcriptional profiling underlined the dependence of hepatic neonatal Tconvs on microbiota signals as we observed an extensive reprogramming between GF and SPF CD44^hi^ Tconvs. In contrast, only minor differences were observed in the Treg compartment (Fig. 1i, Fig. S3h). Activated Tconvs in livers of GF neonates shifted towards a naïve/quiescent transcriptional signature (e.g., *Sell*, *Ccr7*, *Tcf7*) with lower expression of cell-cycle genes (*Mki67*, *Birc5*, *Ube2c*) relative to their SPF counterparts (Fig. 1j). Conversely, activated Tconvs from SPF neonates upregulated transcripts associated with T cell activation (*Il2rb, Id2, Cxcr3*; concomitant with downregulation of *Ccr7, Sell, Tcf7*). Notably however, SPF Tconvs also showed a parallel upregulation of genes encoding inhibitory/tolerance-associated receptors (*Tigit, Pdcd1*) and comprised a higher proportion of CD73^hi^ FR4^+^ anergic cells in flow cytometry (Fig. 1k).

These data fit a model in which the microbiota drives a controlled, tolerance-prone anergy state in Tconvs, seeding an anergic intermediate cell state that can differentiate into peripherally-induced (p)Tregs (*18, 26*). These observations, coupled with the temporal and spatial proximity of liver Tregs and Tconvs in early life, led us to hypothesise that the neonatal Treg wave functions to constrain the Tconv activation that is modulated by the microbiota. In line with this, Ova-specific OTII T cells adoptively transferred into WT neonates showed a significantly decreased proliferation upon antigen exposure in the liver, whereas there was comparable OTII proliferation in liver-draining lymph nodes and other lymphoid tissues in neonates and adults (Figs. S4a-b). Decreased proliferation persisted when antigen was presented endogenously by transgenically modified DCs (cDOG), suggesting an intrinsic immunosuppressive milieu in the neonatal liver rather than insufficient antigen availability and uptake (Fig. S4c). To directly test whether neonatal Tregs exert a suppressive effect on microbiota-induced Tconv activation, we made use of the DEREG mouse model, in which Tregs can be transiently depleted through the administration of diphtheria toxin (DT). Treg depletion during the first week of life strongly amplified hepatic CD4^+^ Tconv activation and frequency (Fig. 1l). Activated T cells upregulated CD69 and CXCR6 together with PD-1, resulting in an aberrant pro-inflammatory phenotype in the absence of Treg-dependent control, reminiscent of auto-aggressive liver T cells (Fig. 1l) (*27*). Collectively, these data demonstrate that the neonatal hepatic Treg wave is developmentally programmed, microbiota-independent, and functions to restrain the microbiota-dependent Tconv activation in the liver.

### The neonatal T cell wave is characterized by clonal proliferation and a shift to an effector phenotype of Treg and Tconv

We next investigated the antigen-dependence and clonal architecture of the early life hepatic T cell wave. Notably, without exposure to cognate antigen, there was no detectable neonatal burst of either Tregs or CD44^hi^ Tconvs in the liver of Ova-specific OTII Rag^−/-^ mice (Extended data 4d). These data led us to hypothesize that both T cell subsets primarily respond to cognate antigen presentation and to examine the clonal repertoire of the hepatic neonatal T cells embedded in the scRNA-seq dataset introduced in Fig. 1 (Fig. S3).

In adult and very young (PND4) animals, the hepatic T cell population was almost exclusively polyclonal (Fig. 2a). In contrast, livers of neonatal mice after PND4 contained a markedly increased proportion of expanded T cell clones, coinciding with the neonatal T cell burst. These findings suggest that the expansion is at least partly clonal and driven by antigen-specific recognition. Interestingly, while there was minimal clonotype sharing between the Treg and the Tconv population in the liver of adult mice, mice at PND7 and older exhibited expanded shared clones, coinciding with the peak of the neonatal T cell expansion and consistent with the possibility that some of the hepatic neonatal Tregs are peripherally induced (Fig. 2b).

**Figure 2:**
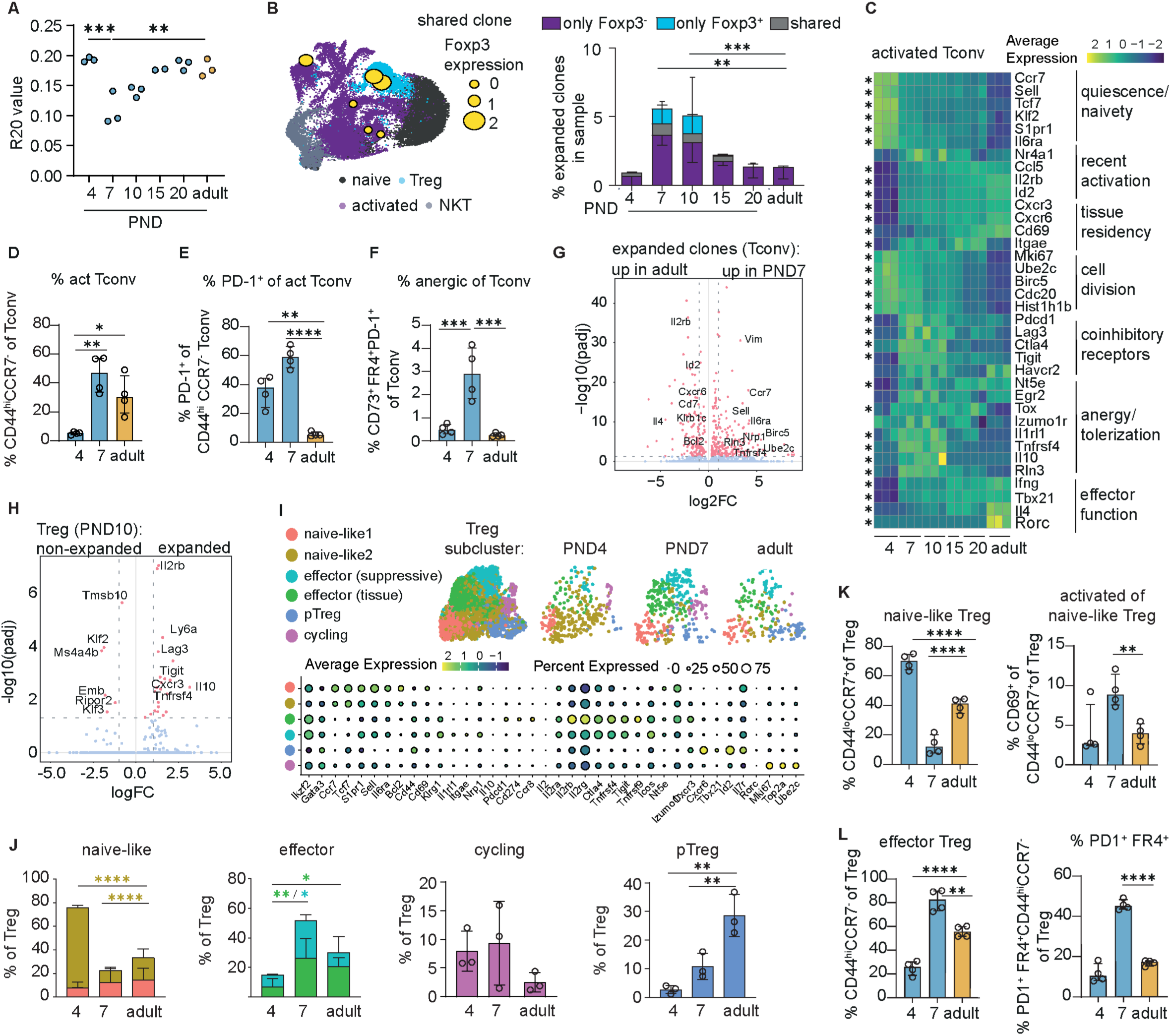
Clonally expanding Tregs and Tconv co-evolve into effector and anergy programs during in neonatal hepatic tissues. (A) R20 score (defined by the fraction of unique clones, in descending order of frequency, which account for 20 % of the sequenced repertoire) of clonal repertoire per mouse from VDJ scRNASeq. 2-3 mice per limepoint. One Way ANOVA + Dunnetrs multiple comparison to adult group. (B) % clones per mouse that are expanded (>1 cell of the same clonotype). 2-3 mice per limepoint composition of the expanded clones (either only Foxp3-cells in a clones purple,) only *Foxp3^+^* (blue) and shared clones (both *Foxp3^+^* and *Foxp3-* cells in one clone). Two Way ANOVA + Dunnetrs multiple comparison to adult group. Representative depiction of a shared clone on a UMAP with cells depicted as yellow circles according to their *Foxp3* expression level). (C) Heatmap of canonical genes expressed over lime in activated Tconv (purple cluster in A). Asterisks indicate DE gene expression between an early neonatal lime point (PND4 or PND7) and adult lime point. (D) % activated (CD44^h1^CCR7^lo^) Tconv measured via flow cytometry, (E) % PD-1^+^ and % anergic (CD44^h1^PD-1^+^ FR4^h1^ CD73^h1^) cells of activated Tconv, (F) % anergic of total Tconvs measured via flow cytometry. 3-4 mice per lime-point from 1 representative experiment; One Way ANOVA + Tukey’s post hoc test. (G) Volcano plot with DE genes between Tconvs from expanded clones (>1 cell of the same clonotype) in 7-day old and adult mice. (H) Volcano plot with DE genes between expanded Foxp3+ cells(>= 2 cells per clone) and non-expanded Foxp3+ cells (<2 cells per clone) at PND10. (I) Reclustering of Tregs (blue cluster in A), bubbleplot displaying cluster marker genes used for identification, and UMAPs depicting composition at different lime points after birth. (J) % of cells in a certain cluster within Tregs at PND4, PND7 and adult. One-Way ANOVA + Tukey’s post hoc test. (K) % CD44^lo^ CCR7^hi^ of Tregs and CD69^+^ of CD44^1^° CCR7^h1^ Tregs measured by flow cytometry and (L) % CD44^h1^CCR7-ofTreg and PD1^+^ FR4^+^ ofTreg measured by flow cytometry. One Way ANOVA + Tukey’s post hoc test. All experiments containing flow cytometry were done with the compared groups within 1 experiment processed on the same day. Asterisks indicate significance levels: * < 0.05, ** < 0.01, *** < 0.001 **** < 0.0001. All tests were performed in a two-sided manner.

Consistent with the clonal repertoire data, activated Tconvs at PND4 displayed expression of naïve markers as well as cell cycle genes, indicating recent activation. By PND7, activated neonatal Tconvs acquired adult-like activation markers but also expressed inhibitory or anergy-associated genes (e.g., *Havcr2*, *Lag3*, *Tigit*, *Tnfrsf18* with reduced *Il2*), consistent with a controlled activation-to-anergy transition (Fig. 2c). Consistently, flow cytometric analysis of surface marker expression revealed an initial activation of Tconvs between PND4 and PND7 (Fig. 2d). Furthermore, a temporary increase of the fraction of PD-1^+^ cells and cells with an anergy-phenotype was observed within activated Tconv at PND7 (Fig. 2e-f). Comparing the phenotype of Tconv clones between neonate and adult revealed upregulated proliferation genes (*Mki67, Birc5, Ube2c*) alongside tolerance-linked regulators (*Tnfrsf4, Rln3, Nrp1*) in neonates, while in adults, the few expanded clones preferentially expressed tissue-resident and effector-associated genes (*Cxcr6, Klrb1c, Il4*; Fig. 2g).

Among neonatal Tregs, expanded clones preferentially expressed *Il10*, *Lag3*, *Tigit*, *Tnfrsf4* (OX40), highlighting a suppressive effector program of clonally engaged Tregs (Fig. 2h) and reflecting the strong transcriptomic changes in neonatal Tregs in the early postnatal period (Fig.s 2i-j). Most hepatic Tregs bore a resting or “naïve-like” signature characterised by expression of *Tcf7, Sell*, *Ccr7* and others at PND4 (Fig.s 2i-j), corresponding to a CD44^lo^ CCR7^+^ surface phenotype that shifted to a more activated phenotype at PND7 (Fig. 2k). The Treg expansion at PND7 was accompanied by a loss of the naïve-like Treg subset and a concomitant transition to an effector-Treg state (CD44^hi^ CCR7-; Fig. 2l) characterized by increased expression of *Pdcd1* (encoding PD-1), *Ctla4* and *Klrg1* (Fig. 2i-j) as well as surface protein expression of FR4 and PD1 (Fig. 2l). At both early time points (PND4, PND7) Tregs also showed increased expression of proliferation markers (*Mki67*, *Top2a*) suggesting that the increasing number of hepatic Tregs is at least partially due to local proliferation (Fig. 2j).

In addition, starting from PND7, a gradual accumulation of a Treg population marked by lower *Ikzf2* and higher *Cxcr6*, *Cxcr3*, *Tbx21* expression could be observed - these Tregs likely are a peripherally derived, tissue-imprinted Treg subset and represent a major population of adult hepatic Tregs, which was also evidenced by a higher proportion of CXCR6-expressing Tregs in adult livers (Fig. 2h-i; Fig. S4e). Finally, only a minor population of Tregs in the liver expressed *Rorc* (encoding RORγt) at any of the time points analysed, suggesting minimal RORγt⁺ pTreg contribution. (Fig. 2i; Fig. S4e).

Together, these data define the neonatal hepatic T cell wave as clonal and antigen-specific comprising both Tregs and Tconvs. Influx of early thymic Tregs led to differentiation of effector-suppressive tissue Tregs that establish a regulatory milieu in the neonatal liver. In parallel, naïve CD4⁺ T cells entering the liver undergo local activation. Interestingly, the clonal overlap suggests some naïve T cell conversion into peripherally induced Tregs, likely with distinct antigen specificities from the main Treg wave. The coexistence of thymic and peripherally induced tissue Tregs may therefore provide layered control over neonatal Tconv activation within the hepatic niche. By adulthood, this layered architecture condensed into a smaller CXCR6⁺ Treg compartment with reduced proliferative activity but maintained tissue residency, suggesting that the broad effector–suppressive spectrum of PND7 Tregs narrows into a stable, specialized pool reflecting the altered immune-functionality of adult hepatic tissue.

### Neonatal hepatic Treg expansion is driven by intrahepatic DC1-mediated antigen presentation

To directly address the mechanisms of induction and action of neonatal hepatic Tregs, we performed an unbiased ligand–receptor screen on the neonatal scRNA-seq kinetic (Fig. S5a-b), which highlighted cell-cell interactions guided by checkpoint axes. Specifically, the PD-L1 and PD-L2 signalling pathways were enriched in neonates, with expression of *Cd274* (PD-L1) on neonatal CCR7⁺ DCs and KCs, *Pdcd1lg2* (PD-L2) on neonatal NKs, ILCs and CCR7^+^ DCs, and strong *Pdcd1* (PD-1) expression on neonatal Tregs (Fig. 3a; Figs. S5c-d). In line with this, PD-L1 inhibition in neonates induced a shift in the regulatory tone: The frequency/number of Tregs increased, whereas Tconv numbers remained unchanged, leading to an increased Treg/Tconv ratio. (Fig. 3b; Figs. S5e).

**Figure 3:**
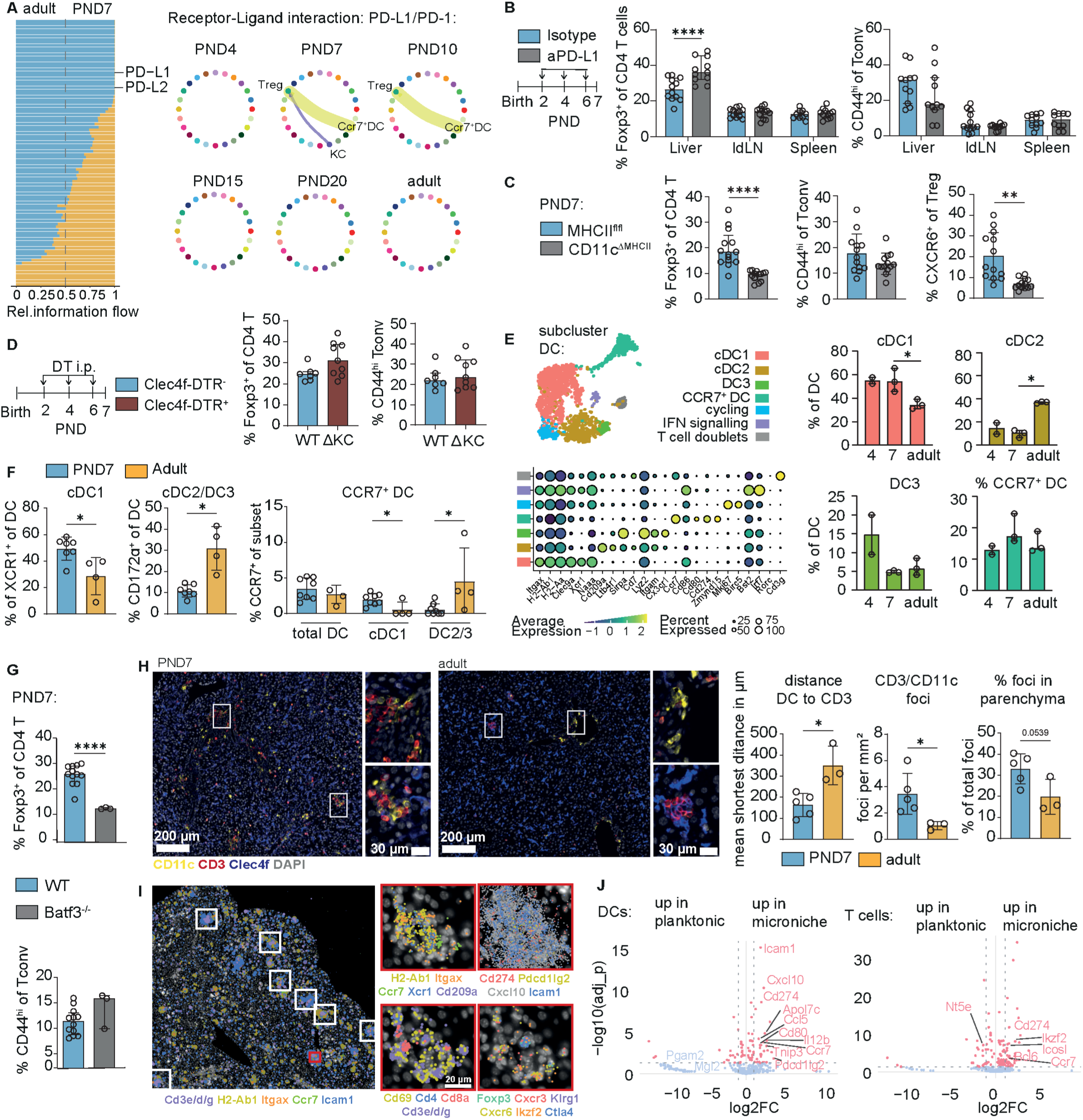
CCR7^+^ cDC1 assemble MHCll-dependent DC-T cell microclusters that enforce PD-L1/PD-L2-tuned tolerance. (A) Overview of differentially expressed ligand receptor pairs predicted by CellChalDB between PND7 and adult CD45^+^ cells and circle plots displaying kinetic overview of predicted POL1-PD1 interaction of the Treg cluster with other cell populations. Circles represent clusters/subsets from UMAP in Ext. Data 3c. (B) Experimental design for aPD-L1 blockade in neonatal WT mice and flow cytometry data of % of Treg within CD4 T cells as well as % of CD44^hi^ within Tconv in liver, ldLN and spleen. Multiple Mann Whitney U tests. (C) Flow cytometry data of % of Treg within CD4 T cells as well as % of CD44^hi^ within Tconv in liver in CD11cCre/+MHCllflfl mice (CD11 cL’:.MHCII) and their CD11c+l+MHCII” (MHCll”)littermates. Mann Whitney U tests. (D) Experimental design for Kupffer cell (KC) depletion in neonatal Clec4fDTR mice and their WT litter mates; flow cytometry data of % ofTreg within CD4 T cells as well as % of CD44^hi^ within Tconv in liver. Mann Whitney U tests. (E) scRNASeq reclustering of dendritic cells from PND4 to adult hood with canonical marker genes displayed in bubbleplot and kinetics of the subclusters. One Way ANOVA + Tukey’s comparison. (F) Flow cytometry comparison of the liver DC compartment of neonatal WT and adult mice. Mann Whitney U tests. (G) Flow cytometry data of % of Treg within CD4 T cells as well as % of CD44^hi^ within Tconv in liver of neonatal WT and BATF3_,_ mice at PND7, liver Treg frequency and CD44^hi^ Tconv frequency. Welch I-tests. (H) Multiplex immunofluorescence images and quantitative analysis of whole tissue slides; mean shortest distance between T cell (CD3+) and closest DC (CD11 c^+^Clec4flba1 t density of manually counted CD3/CD11 c foci in tissue and % of foci identified as in parenchyma by absence of vessels in close proximity. Welch I-tests. (I) Identification and characterization of microclusters in Xenium targeted spatial transcriptomics of neonatal liver (PND7). Left pictures shows indicated gene expression + T cell-DC clusters found by DBScan analysis highlighted in rectangles. Close-ups show the red highlighted cluster and expression of several transcripts of interest. (J) DE gene analysis of DCs and T cells inside and outside of microniches from Xenium panel. Top DE genes of cell type specific probes highlighted with labels. All experiments containing flow cytometry were done with the compared groups processed on the same day within 1 experiment (except BATF3-/- and WT pups). Asterisks indicate significance levels: * < 0.05, ** < 0.01, *** < 0.001 **** < 0.0001. All tests were performed in a two-sided manner.

Since our results demonstrated that neonatal Treg expansion was antigen-dependent (Fig. S5d), we hypothesised that either hepatic CCR7^+^ DCs or KCs act as principal APCs. To analyse their relative contribution, we made use of CD11c-Cre×H2-Ab1^fl/fl^ mice, which lack MHCII expression on DCs as well as Clec4f-DTR mice, in which KCs can be depleted. At PND7, a DC-specific MHCII deficiency resulted in a significant reduction in the Treg frequency as well as reduced Treg CXCR6 expression. Notably, the number of activated Tconvs was unchanged (Fig. 3c; Fig. S6a). In contrast, depletion of Kupffer cells or the deletion of MHCII on hepatocytes did not affect Treg frequencies nor Tconv activation status (Fig. 3d; Figs. S6b-c). Collectively, these results identify antigen presentation by DCs as the principal mechanism of neonatal hepatic Treg induction.

Transcriptome and flow-cytometry analyses identified a predominance of the cDC1 subpopulation in the neonatal liver, whereas the proportion of cDC2/DC3s increased after the postnatal period (Fig. 3e-f; Figs. S6d-g). Notably, despite its role in tolerance induction in the neonatal intestine(*28, 29*), we could not detect the presence of the tolerogenic RORγt⁺ DC subset in the neonatal liver (Fig. 3e). While the overall fraction of CCR7⁺ DCs remained unchanged between adults and neonates, the majority of the CCR7⁺ DC pool shifted from cDC1s in neonates to cDC2/DC3s in adults (Fig. 3f) suggesting that cDC1s may act as the principal APCs regulating the hepatic neonatal Treg wave. The decreased frequency of hepatic Tregs in cDC1-deficient BATF3 KO pups corroborated the role of cDC1s in regulating Tregs in the neonatal liver tissue (Fig. 3g). Crucially, postnatal liver T cell accumulation did not depend on migration of DCs to the LNs, as T cell density was not reduced in neonatal livers of CCR7^−/−^ mice (Fig. S7a).

In line with this, multiplex immunofluorescence microscopy revealed that T cells in neonatal liver tissue were organised in microclusters and colocalised with CD11c⁺ Iba1⁻ Clec4f⁻ DCs as well as Tconvs (Fig. 3h). These DC-T cell microclusters were significantly more prevalent in neonatal compared to adult livers. Moreover, they had a distinct localisation: in adult tissues the DC-T microclusters were mainly confined to periportal areas, while they were abundant in the liver parenchyma of neonates (Fig. 3h; Fig. S7). Crucially, spatial transcriptomics confirmed co-localization of DCs, Treg and Tconv in microclusters in the neonatal liver (Fig. 3i; Figs. S7 a-e). DCs in neonatal liver microniches expressed significantly more *Ccr7, CD80, Pdcd1-lg2, Cd274* corresponding to CCR7^+^ DCs (Fig. 3j; Fig. S7f), while T cells showed a significant enrichment of Helios (*Ikzf2* transcript) within the clusters (Fig. 3j; Figs. S7g-i). Notably, DC-T cell clusters in neonatal livers showed a high level of *Cxcr3* transcripts and its ligand *Cxcl10* suggesting a chemokine-dependent origin of these cell clusters (Fig. 3i; Figs. S7 d-e).

Together, the presented results identify anatomically discrete, cDC1-anchored regulatory clusters in the neonatal liver that (i) depend on MHCII on CD11c⁺ DC, (ii) assemble CCR7⁺ cDC1–Treg–Tconv triads in parenchyma and near portal tracts, and (iii) use PD-L1/PD-L2 checkpoints to stabilize tolerance as the liver transitions postnatally. These data align with postnatal portal-area tissue remodelling and the zone-specific maturation of hepatic stromal compartments, providing a structural basis for immune organisation in early life (*30, 31*).

### The neonatal liver environment confers tolerance to *in situ* activated T cells with consequences for infection

We next asked whether this strong immunoregulatory environment in the neonatal liver tissue may be detrimental during hepatotropic infection. Using the Norway rat hepacivirus (NrHV) model (a mouse HCV surrogate (*32*)), we infected PND7 neonates and adults. Despite the higher baseline density of Tconv and CD8^+^ T cells in neonatal liver tissues (see Fig. 1), viral clearance was delayed in neonates relative to adults (Figs. 4a-b; Fig. S8a). Seven days post-infection, neonates displayed a robust, liver-specific induction of Tregs, which was not seen in adult mice. Neonatal Tregs also showed increased expression of effector and suppressive markers (Figs. 4c-d; Fig. S8b). Nevertheless, neonatal Tconv and CD8^+^ T cell densities in the neonatal livers increased strongly during the course of infection, and upregulated activation markers (Figs. S8c-d). However, Tconvs showed decreased effector marker expression in infected neonatal mice when compared to adult infected mice (Fig. S8c), while CD8^+^ T cells were activated to the same level in neonatal and adult infected mice (Fig. S8d). To test whether neonatal Tregs causally influenced the antiviral host response kinetic, we transiently depleted Tregs in neonates shortly after infection (Figs. 4f-g) using a previously established protocol to induce virus clearance in chronically infected adult mice (*33*). Treg depletion led to a decreased viral load and accelerated viral clearance, consistent with a functional role of the developmental Treg wave in modulating early antiviral immunity (Figs. 4f-g). Taken together, the neonatal hepatic niche enforces tolerance on freshly activated T cells, beneficial under commensal colonization to prevent immunopathology (*17, 18*), yet detrimental during hepatotropic virus infection, where it tempers effector function and delays viral clearance.

**Figure 4:**
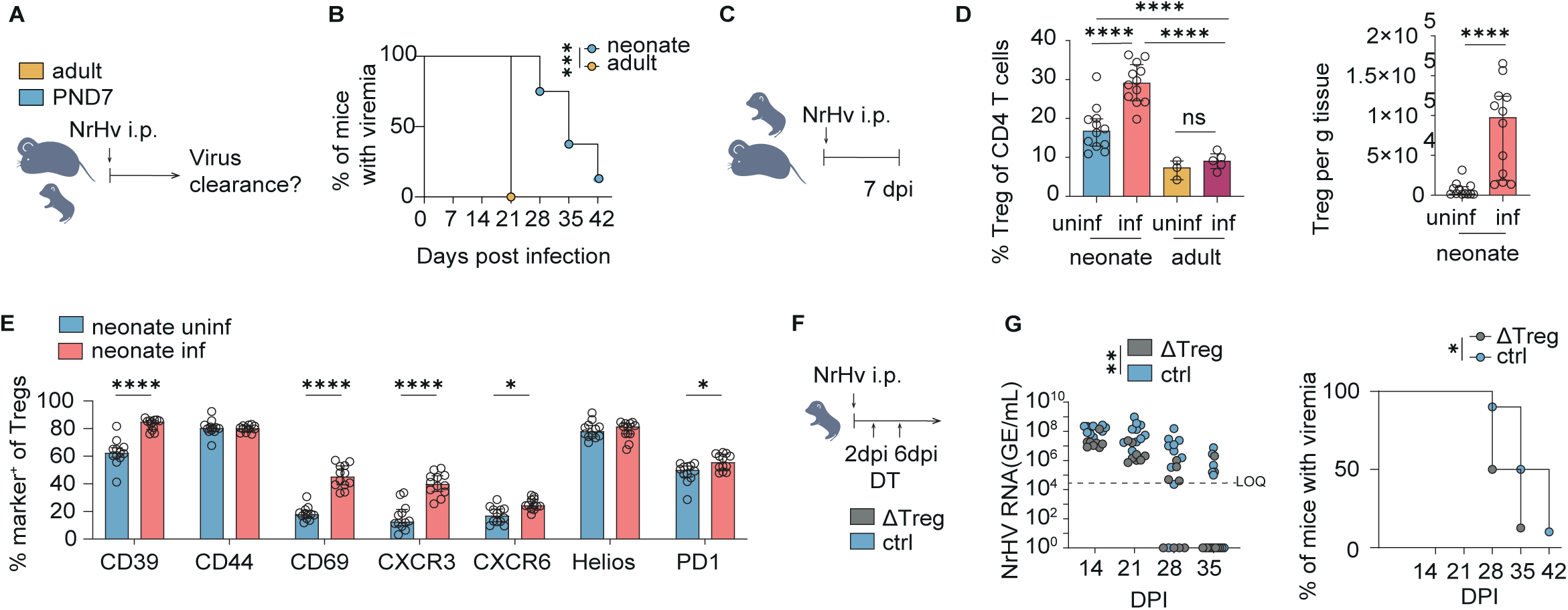
Tolerance-biased neonatal liver delays viral clearance in a hepatitis C model despite robust T-cell activation. (A) Experimental design of NrHV infection model. (B) Clearance rates of neonatal mice infected at PND7 and infected adult mice. 5-10 mice from 1 adult and 2 independent neonatal experiments. Kaplan Meier survival curve + log-rank test. (C) Experimental design of NrHV infection model for acute infection flow cytometry analysis. (D) % Treg of CD4 T cells in liver tissue of uninfected and infected neonates and adults 7 days post infection (dpi) and Treg density in liver tissue of infected and uninfected neonatal mice. Flow cytometry data of 11-12 neonatal mice from two independent experiments and 3-5 adult mice from 1 experiment. One Way ANOVA + Holm-Sidak multiple comparison tests. Treg density: Mann Whitney U test. (E) Phenotypic markers of liver Treg in neonatal uninfected and infected and adult infected mice 7 days post-infection measured by flow cytometry. 12 neonatal mice per group from 2 independent experiments. Multiple Mann Whitney U tests + Holm-Sidak multiple comparison correction. (F) Experimental design of Treg depletion during NrHV infection in neonates. (G) Viremia levels in NrHV infected Foxp3-DTR mice receiving 2 doses of diphtheria toxin (DT) and infected WT control neonates 2 and 6 dpi. 8-10 neonatal mice per group pooled from 1-2 independent experiments. Two Way ANOVA (blood viremia levels) and Kaplan Meier survival curve +log-rank test. Asterisks indicate significance levels: * < 0.05, ** < 0.01, *** < 0.001 **** < 0.0001. All tests were performed in a two-sided manner.

### Early-life Treg programs shape adult susceptibility to diet-induced Metabolic Dysfunction-Associated Steatotic Liver Disease (MASLD)

To test whether early-life Tregs seed long-lived regulatory compartments, we lineage-tagged CD4⁺ T cells either in neonates or adults using CD4-CreERT2;Rosa-YFP mice and analyzed them 8 weeks later (Fig. 5a). While there was no difference in the overall proportion of YFP^+^ CD4^+^ T cells in the liver (Fig. S9a), neonatal timestamping yielded a disproportionately high fraction of YFP⁺ Tregs in adult liver, whereas adult timestamping contributed minimally to the hepatic Treg pool (Fig. 5b). Comparing YFP⁺ vs. YFP⁻ compartments showed that early-life-tagged CD4⁺ cells were strongly biased toward a Treg fate in the liver, consistent with (i) long-term retention of neonatal thymic Tregs and/or (ii) preferential pTreg differentiation from tissue-imprinted effectors within the neonatal hepatic niche (Fig. 5c). We next asked whether these neonatally generated persisting Tregs had lasting consequences. For this, we timestamped neonatal CD4^+^ T cells and challenged the mice as adults with a choline-deficient, high-fat diet (HFD) that drives steatosis and metabolism-associated steatohepatitis (MASH) over 16 weeks (Fig. S9b). In this model, subsets of effector Tregs contribute to fibrogenic programs under metabolic stress (*34*). Across groups, high-fat feeding increased the overall Treg share in liver CD4^+^ T cells and selectively expanded Helios⁺ Tregs approximately 3-fold. Lineage tracing showed that this expansion also occurred within neonatally derived (YFP⁺) cells, indicating that early-life–imprinted Tregs remain responsive in adulthood (Fig. S9b). Finally, to definitively determine whether the neonatal Treg wave had functional relevance for the development of metabolic liver disease in later life, DEREG neonates were transiently depleted of Tregs during the first postnatal week and subjected to HFD for 16 weeks as adults (Fig. 5d). Neonatally depleted males showed a modest but significant excess weight gain, trending alanine aminotransferase (ALT) levels, and increased histological steatosis, while glucose tolerance was unchanged (Fig. 5e-g; Fig. S9c). Female mice displayed a MASLD phenotype that was independent of neonatal Treg depletion. (Sig. S9d). The pathological MASLD phenotype in neonatally depleted males was accompanied by increased levels of Th1 and Th17 effector cells, indicating a pro-inflammatory skew (Fig. 5h). Notably, the Helios⁺ Treg fraction increased in the HFD group, whereas the total Treg frequency within CD4^+^ T cells was unchanged between the DEREG and WT HFD groups, suggesting an altered Treg composition rather than an absolute loss after early-life Treg depletion (Fig. 5i). Taken together, neonatal Treg programs durably shape the adult hepatic T cell compartment and modulate the susceptibility to metabolic liver disease. Early-life generated T cells preferentially populate the adult liver Treg niche, transient neonatal Treg loss primes male-biased hepatic steato-inflammation under dietary stress, and dietary challenge amplifies a Helios⁺ Treg subset that includes neonatally derived cells.

**Figure 5:**
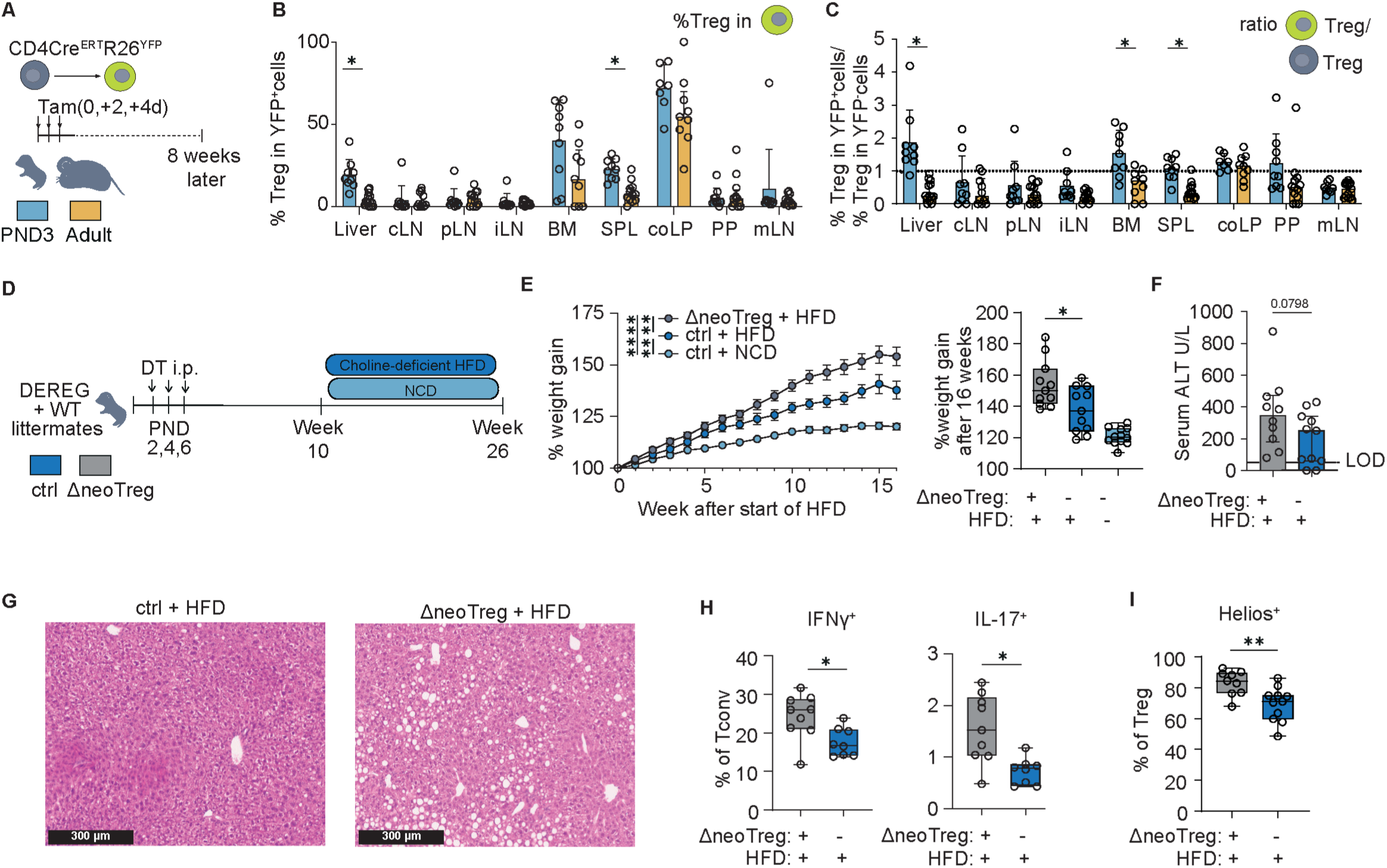
Early-life Treg programs shape adult susceptibility to diet-induced MASLD. (A) Experimental design for time-stamping experiment of T cells in co4c,.rnr R26YFP_model. (B) % of Treg in YFP^+^ CO4 T cells 8 weeks after limestamping per organ and (C) ratio of Treg in YFP^+^ and YFP-CO4 T cells. Mann Whitney U tests. (D) Experimental design of diet induced MASLO experiment with neonatal Treg depletion in OEREG mice and WT littermates. 8-11 mice from each group from 5 independent experiments. Only litters containing both male OEREG and WT littermates were used for HFO analysis. (E) % weight gain during the HFO treatment in neonatally transiently Treg depleted male mice, their littermates and normal chow diet (NCO) fed control mice from (d) (parametric, paired mixed-effects analysis + Holm Sidak multiple test correction) and % weight gain at the end of the experiment (Unpaired I-test). (F) Serum ALT levels in male mice from (d). Unpaired I-test. (G) representative H&E stainings of liver tissue from (H) IFNy^+^ and IL-17^+^ ofTcaonv after restimulation with PMA/lonomycin and (I) % of Helios^+^cell within Tregs measured by flow cytometry. All experiments containing flow cytometry were done with the compared groups processed on the same day within 1 experiment. Asterisks indicate significance levels: * < 0.05, ** < 0.01, *** < 0.001 **** < 0.0001. All tests were performed in a two-sided manner.

## Discussion

Our data identify a disproportionately large CD4 T cell pool in the neonatal liver, compared with other organs, characterized by a pronounced Treg wave that is independent of the microbiota, and a concurrent accumulation of activated Tconv modulated by microbial exposure. TCR/clonotype analyses and transcriptomic/phenotypic characterization of Tregs in early life hepatic tissue point to a biphasic program: an early, developmentally hard-wired tTreg wave that arrives in a naïve-like state and differentiates *in situ* into tissue effector Tregs, followed by a secondary enrichment of peripherally induced Tregs consistent with an anergy-to-pTreg maturation trajectory(*35, 36*). Conceptually, the neonatal hepatic Treg wave fits the broader pattern of early-life regulatory responses at barrier sites. In lung and skin, microbiota-dependent Treg accumulations arise later in the pre-weaning period (around postnatal day 8 in lung and day 13 in skin, respectively) and have been suggested to restrain local inflammatory priming(*14, 15*). In contrast, the neonatal liver hosts an earlier, microbiota-independent expansion, suggesting a developmentally hard-wired regulatory program that precedes and complements microbiota-dependent regulation at barrier tissues. In our data, neonatal Tconv responsiveness tracks the level of microbial exposure, whereas the Treg peak is exposure-insensitive; we interpret this as a division of labor, in which endogenous/self-antigen preferentially sustains Tregs while microbe-associated signals “titrate” Tconv activation. A developmentally programmed, microbiota-independent Treg wave may be essential to calibrate the transition to highly variable postnatal exposure to microbial and dietary antigens.

Functionally, this program maximizes regulatory bandwidth at a time when the peripheral CD4⁺ pool is dominated by recent thymic emigrants (RTEs), a population more self-reactive and less stringently tuned than adult naïve T cells(*1–4*). Within this context, the successive Treg waves likely differ in TCR specificity and function: early tissue Tregs provide rapid local control of Tconv activation as polyreactive RTE-Tregs and non-Tregs first enter the periphery, whereas later Tregs, of either thymic or peripheral origin, consolidate longer-term protection from immune-mediated pathology, including metabolic liver disease.

Where first antigen encounters occur is pivotal: During suckling stage, luminal antigen access to the immune system of the gut is restricted(*10, 13*), yet the portal circulation can still ferry self-derived metabolites and microbiota-derived cues to the liver, in addition to abundant self-derived signals generated by the metabolically active parenchyma. At present, we cannot distinguish whether Tconv tuning is driven primarily by cognate microbial antigens or by non-antigenic PAMPs. A parsimonious explanation is that immature enterocytes engage in bulk endocytosis/macropinocytosis, allowing microbial material, including both antigens and PAMPs, to reach the portal circulation despite reduced classical macromolecule uptake, a process often discussed under the umbrella of “gut closure”(*37, 38*). Within this framework, our data characterize a neonatal hepatic environment in which antigen presentation by cDC1s results in a tolerance-promoting microenvironment along the gut–liver axis, consistent with CD4⁺ T cell hyporesponsiveness and the inhibition of self-reactivity(*36, 39*).

At the cellular wiring level, our imaging and perturbation experiments show that CCR7⁺ cDC1 nucleate APC–T cell clusters that act as tolerogenic foci in the neonatal liver. Within these sites, PD-L1/PD-L2 checkpoints constrain Tconv activation while permitting robust Treg accumulation. Anti-PD-L1 treatment therefore results in selective Treg accumulation without unleashing Tconvs, consistent with differential checkpoint sensitivity of the two compartments rather than a single dominant effector mechanism. This aligns with the broader view that Treg control emerges from precise positioning and multiple, context-dependent mechanisms rather than one canonical pathway(*40*). Notably, while RORγt⁺ DC populations are pivotal in intestinal tolerance, we do not detect a RORγt⁺ DC contribution in the neonatal liver; instead, cDC1 dominate the activated CCR7⁺ pool. This organ-specificity fits with current syntheses that place RORγt⁺ DCs at the center of gut tolerance but not in every tolerogenic niche(*41*).

The anatomical microenvironment of these interactions is unique for hepatic tissue in early life. We observe cDC1-anchored T cell microniches not only in periportal areas but also widely distributed across the parenchyma, an anatomical arrangement that is lost in adulthood. Work on neonatal liver development supports a window in which immune and stromal programs are broadly distributed across the parenchyma and remodel toward the adult architecture around weaning, providing a structural rationale for elevated CD4⁺ density and Treg skew in neonates(*31, 42, 43*). Within this setting, spatial transcriptomics reveals that neonatal DC-T cell clusters are enriched for transcripts associated with leukocyte migration, including *Ccr7*, *Cxcr3* and *Cxcl10*, and colocalize with *Icam1*⁺*Cxcl10*⁺ structural cells that are distinct from hepatocytes. These features support a chemokine- and adhesion molecule-dependent origin of the microniches and resemble Th1 response-enhancing CXCL10⁺ activation niches described in other tissues(*44, 45*). At the same time, their dispersed, T cell–dominated architecture and lack of B cell organization distinguish them from the B cell-rich tertiary lymphoid structures that can arise in chronically inflamed livers and other organs(*21, 23, 24*).

Within this framework, the balanced activation we observe, characterised by clear CD4⁺/CD8⁺ responses to microbial exposure but slower pathogen control under hepatotropic virus challenge, fits a tolerance-biased set point in neonates that prioritizes tissue protection while accepting the trade-off of delayed pathogen clearance(*46*). Transient disruption of neonatal regulation imprints long-lasting vulnerability patterns in adulthood, highlighting its non-redundant functionality and suggesting that the composition and education of the neonatal T cell pool, more than sheer numbers, forecast later liver inflammatory risk(*47*). Paediatric livers exhibit strong tolerogenic features, and early-life hepatotropic virus infections more readily become chronic. Our data provide a mechanistic basis for neonatal hepatic tolerance: a CCR7⁺ cDC1-anchored, regulatory immune architecture that skews activation outcomes and Treg differentiation along the gut–liver axis(*36, 47*). In humans, this may offer a plausible explanation for the age-dependent pathology of hepatitis B virus: newborns typically develop chronic infection, whereas adults more often mount acute, self-limiting disease, consistent with a neonatal liver environment that dampens Tconv/CD8 priming and cytotoxic function, favoring viral persistence.

Notably, in chronic metabolic disease the Treg program can become maladaptive. In mouse MASH, hepatic Tregs expand and adopt an activated CCR8⁺/ST2⁺/KLRG1⁺ state that promotes amphiregulin-dependent stellate-cell activation and fibrosis(*34*). Although the CCR8⁺/ST2⁺/KLRG1⁺ Tregs reported in MASLD/MASH display hallmarks of tissue Tregs, that overlap with features we also observe in neonates, our findings indicate that neonatally derived tissue Tregs are protective in the long term. Tissue Treg pools in non-lymphoid organs, including in the liver, are now understood to be shaped by transient multi-tissue migration and a conserved residency program rather than life-long persistence of a fixed neonatal cohort(*47*).

Two non-exclusive factors may reconcile this with reports of pathogenic CCR8⁺/ST2⁺/KLRG1⁺ Tregs in established disease. First, kinetics: neonatal tissue Tregs are in place at baseline, shaping responses from the onset, whereas MASLD-associated Tregs are recruited or reprogrammed during disease and may act at later, injury-rich timepoints, where their repair/EGF-family programs can inadvertently foster fibrosis. Second, TCR specificity: neonatal tissue Tregs are likely selected to distinct antigenic landscapes (self, perinatal antigens) that differ from the damage- and lipotoxicity-derived antigens encountered in MASLD. Thus, despite similar surface programs, their TCR repertoires and instructive cues may drive divergent functions. This framework predicts that early-life Treg ontogeny/timing and TCR repertoire, rather than phenotype alone, will stratify MASLD risk and outcomes.

## Acknowledgements

We would like to thank Jochen Huehn (Helmholtz Centre for Infection Research) for critical discussion, Dr. Martin Guilliams and Dr. Alain Beschin (VIB, Belgium) for providing the Clec4F-DTR mouse line under a material transfer agreement with RWTH Aachen University, Dr. A. Viehof, J. Bosch and A. Panyot (University Hospital of RWTH Aachen, Functional Microbiome Research Group) for support with gnotobiotic mice and Dr. Marijana Basic and Dr. Andre Bleich (Hannover Medical School) for supply of germ-free mice, Dr. J. Wang for help with gnotobiotic animal handling (Bern University), Nelson Gekara and Kyaw Min Aung (Freiburg University Hospital and Umea University) for the generation of *E.coli* MVs and the histology, genomics and flow cytometry core facilities (German Research Foundation (DFG) grant project ID 439895892) of the Interdisciplinary Center for Clinical Research (IZKF) Aachen within the Faculty of Medicine at RWTH Aachen University. We also would like to thank the society of mucosal immunology for supporting ELS with a technique sharing grant to perform the reversible colonisation experiments at DBMR Bern. Language editing assistance was provided using ChatGPT (OpenAI, V4.5 and V5). The authors verified all edits.

## Funding

Federal Ministry of Research, Technology and Space (BMFTR) MICROSTAR fund to (NT)

RWTH Aachen Faculty of Medicine START-Program (ELS, NT)

German Research Foundation (DFG), Collaborative Research Center CRC1382, grant 403224013 (MWH, TC, OP, AG and NT)

German Research Foundation (DFG), Clinical Research Unit 344, grant 417911533 (NT)

German Research Foundation (DFG), TRR 359 PILOT, grant 491676693 (SPR, TC, MWH and NT)

European Research Council (ERC), grant 101019157 (MWH)

National Institutes of Health (NIH), grant R01AI170725 (EB)

Stiftung Molekulare Biomedizin, Peter Hans Hofschneider Professorship (SCVG)

Swiss National Science Foundation (SNSF), grant 310030_21251 (SCVG).

## Author contributions

Conceptualization ELS, EB, MWH, NT; Methodology ELS, THN, DM, MK, ALG, SPR, SGV, NT; Software THN, ALG, AG, Validation ELS, NT; Formal Analysis ELS, THN, MK, ALG, AG; Investigation ELS, DM, MK, JP, SAB, JB, YC, MU, CK, JLS, JH, ASS; Resource ALG, ChK, DV, SPR, OP, TC, AG, EB, MWH, NT; Data curation ELS, THN, DM, MK, ALG, AG; Writing-original draft ELS, VC, NT; Writing-review editing ELS, VC, EB, TC, MWH, NT; Visualization ELS, THN, ALG; Supervision MWH, NT; Project administration ELS, NT; Funding acquisition ELS, MWH , TC, NT.

## Data availability

scRNASeq raw FASTQ data has been deposited in ENA with accession code PRJEB104713. The processed data is available in Zenodo (DOI:10.5281/zenodo.17940609, available from 31/12/2025).

10X Genomics Xenium raw and processed data have been deposited in Zenodo (DOI:10.5281/zenodo.17940609).

## Supplementary files

### Methods

All animal experiments were performed in compliance with the German animal protection law (TierSchG) and approved by the local animal welfare committees, the Landesamt für Verbraucherschutz und Ernährung (LAVE), North Rhine Westfalia (M2025-728, 81-02.04.2017, A460; 81-02.04.2018.A386), Lower Franconia Government and Regierungspräsidium Freiburg, Freiburg (X-20/05F), the Cantonal Veterinary Commission of the Canton of Bern, Switzerland (BE-132/23). Viral infection experiments were performed under specific pathogen-free conditions at the Institute for Animal Studies of Albert Einstein College of Medicine using C57BL/6 or FOXP3-DTR mice (Jackson Laboratory). All mouse experiments were conducted in accordance with the NIH Guide for the Care and Use of Laboratory Animals and approved by the Albert Einstein College of Medicine Institutional Animal Care and Use Committee (protocol number 00001108).

C57BL/6J wild-type (WT), B6.Cg-Tg(TcraTcrb)425Cbn Rag1^tm1Mom/J^ (OTII Rag-/-), C57BL/6-Tg(Foxp3-HBEGF/EGFP)23.2Spar/Mmjax (DEREG), B6-Clec4^ftm1Ciphe^ (Clec4f-DTR) mice, B6.129S(C)-*Batf3^tm1Kmm^*/J, Tg(Itgax-cre)1-1Reiz x H2-Ab1^b-tm1Koni^ , Tg(Cd4-cre/ERT2)11Gnri x B6.129X1-*Gt(ROSA)26Sor^tm1(EYFP)Cos^*/J , Tg(Alb1-cre)7Gsc x H2-Ab1^b-tm1Koni^ , CCR7^gfp/gfp^ (CCR7 KO) were bred locally and held under specific pathogen–free (SPF) or germ-free (GF) conditions at the Institute of Laboratory Animal Science at RWTH Aachen University Hospital, or Institute for Hygiene and Microbiology Würzburg. C57BL/6NTac wildlings were bred locally at the animal facility of the Medical Center – University of Freiburg, Germany. C57BL/6NTac murine pathogen free (MPF) control mice were originally purchased from Taconic Biosciences, subsequently bred locally and housed under SPF conditions at the animal facility of the Medical Center - University of Freiburg, Germany. Wildlings and C57BL/6NTac MPF mice were age matched for all experiments.

### In vivo models

#### Perinatal Treatment with Broad-spectrum Antibiotics

Pregnant dams were treated from E14.5 to PND7 of the offspring with ampicillin (1 mg/mL), neomycin (0.5 mg/mL), vancomycin (1 mg/mL) ad libitum in the drinking water. Additionally, dams were gavaged i.g. once per day with 0.2 mg/g body weight metronidazole dissolved in drinking water for depletion of anaerobic bacteria as metronidazole could not be delivered ad libitum due to the bitter taste of this antibiotic(*48*). The antibiotic-containing drinking water was prepared and exchanged daily.

#### Reversible Colonisation with E. coli HA107

Germ-free mice were either colonised during pregnancy (gestational colonisation of the dam; GC) or in the first week after birth at PND3 and PND5 (Postnatal colonisation of the pups; PC) the E. coli strain HA107 that is auxotrophic for the amino acids meso-diaminopimelic acid and D-alanine(*49*). HA107 was grown in 200 mL LB medium containing 100 µg/mL meso-diaminopimelic acid and 400 µg/mL D-alanine overnight at 37 °C on a shaking incubator and washed twice in sterile PBS. For gestational reversible colonisation, pregnant germ-free dams were gavaged i.g in gnotobiotic isolators as previously described(*50*) at E7, E9, E11 and E13 of the pregnancy with 10^10^ CFU. For postnatal reversible colonisation, germ-free pups were colonised with 7×10^8^ CFU HA107 at PND3 and 10^9^ at PND5 by gavage i.g. within a sterile environment under the laminar flow hood.

#### E.coli outer membrane vesicles (OMV) Treatment

E. coli OMV were prepared by the lab of Nelson Gekara at Umea University, Sweden as described by(*51*). OMVs were administered i.g. (25 mg in 20 µL per pup) to 10-day old pups in PBS eight hours before analysis.

#### Treg depletion

Diphtheria toxin (Merck, No. 322336) was used to deplete regulatory T cells in neonatal DEREG mice. DEREG and WT mice were injected intraperitoneally with 25, 50 and 75 ng DT respectively on PND (Postnatal day) 2, PND4 and PND6 in 10-20 µL PBS.

#### Antibody Treatments

*α*PDL1 blockade antibody (Clone: 10F.2H11; InVivoMAb, Bioxcell) was applied to WT pups at PND2, PND4 and PND6 i.p. 10-20 µL in sterile PBS. Littermates were treated with isotype-matched control antibody (InVivoMAb rat IgG2b isotype control, *α*-keyhole limpet hemocyanin, Bioxcell).

#### KC depletion

Diphtheria toxin (Merck, No. 322336) was used to deplete regulatory T cells in neonatal B6-Clec4ftm1Ciphe (Clec4f-DTR) mice. DTR+ and DTR-mice were injected intraperitoneally with 25, 50 and 75 ng DT respectively on PND (Postnatal day) 2, PND4 and PND6 in 10-20 µL PBS.

#### NrHV infection

Norway rat hepacivirus (NrHV) stocks were generated from a cDNA clone as described previously(*32, 52*). Adult C57BL/6 mice (8–10 weeks) were infected intravenously via the retro-orbital sinus; neonatal mice were infected intraperitoneally at postnatal day (PND) 7 with 10⁴ genome equivalents (GE) NrHV in sterile PBS. Serum was collected at the indicated time points and viral RNA copies were quantified by RT–qPCR targeting the NrHV NS3 region using an in-vitro-transcribed RNA standard curve, essentially as described(*32*).

For Treg depletion during infection, neonatal FOXP3-DTR and WT littermate controls received diphtheria toxin (25 ng/g body weight in 50 µL PBS, i.p.) at 2 and 6 days post infection. Mice were sacrificed at 7 days post-infection. Livers and spleens were collected into 10 mL and 7 mL of RPMI supplemented with 10 % FCS, T cells were isolated immediately. Tissues were minced and digested in HBSS-based digestion buffer containing 0.01 % collagenase IV, 40 mM HEPES, 2 mM CaCl₂, and 2 U/mL DNase I for 12–15 min at 37°C, followed by mechanical homogenization. Subsequent lymphocyte isolation was performed according by density gradient according to “liver cell isolation”. Viability staining was performed first using the Zombie NIR Fixable Viability Kit (BioLegend). Surface staining was conducted at 20°C for 15 min. Intracellular staining was performed following permeabilization using the FOXP3/Transcription Factor Staining Buffer Set (eBioscience), with a 20 min permeabilization at 4°C followed by a 30 min intracellular stain at 4°C. After staining, cells were fixed in 4 % paraformaldehyde (PFA).

#### Time Stamping

50 µg/g body weight tamoxifen (Sigma) was applied three times in 7.5 µg/µL in Clinoleic Solution (Baxter) either to 8-12 week old mice or to neonatal mice on PND3, PND5 and PND7 after birth by intragastric gavage.

#### Choline-Deficient High-Fat Diet

For induction of metabolic dysfunction associated with steatotic liver disease after the different neonatal interventions, mice were placed on a choline-deficient high-fat diet (HFD; (cat. no. D05010402; Research Diets) at the age of 10 weeks for a period of four months. On the last day of the dietary treatment, oral glucose tolerance was tested by fasting the mice for 6 h. Baseline glucose levels were measured with a standard blood glucose monitoring device (AccuCheck) from a drop of tail vein blood. Mice were then gavaged i.g. with glucose (2 mg/g body weight) dissolved in drinking water. Blood glucose levels were measured after 15 min, 30 min, 60 min and 120 min.

#### Isolation of liver immune cells

Livers were perfused with PBS/3 % FCS via the vena cava; the gall bladder was removed, and the liver was weighed and cut into pieces. Liver pieces were either digested in 10 mL of RPMI with Liberase*^TM^*(Roche; 30 µg enzyme/mL digestion solution) and DNase (Roche; 10 µg enzyme/mL digestion solution) for 30 min at 37 °C in a shaking water bath. Tissue remnants were homogenised by processing through a 10 mL syringe, filtered through a 100 µM cell strainer, and rinsed with 10 mL PBS/FCS to retain a single-cell suspension, which was centrifuged for 8 min at 400*×* g to pellet the harvested cells. In some experiments liver cells were isolated by mechanical dissociation (for scRNASeq and in case of CD62L flow cytometry staining): up to 400 mg of liver tissue was placed in the chamber of the Tissue Grinder device with 800 µL of PBS/3 % FCS (FFX Technologies), and mechanically dissociated using the Liver Program of the device twice; the tissue grinder tube was then centrifuged for 5 min at 400*×* g to pellet the harvested cells.

The obtained pellet from both methods was then resuspended in 4 mL of a 40 % solution of phosphate-buffered PBS/Percoll polymer in RPMI/FCS and carefully layered over 4 mL of a 70 % solution of phosphate-buffered Percoll polymer (Cytiva). Density gradient centrifugation was performed for 25 min at room temperature at 700*×* g without brake and acceleration set to level 3. After the gradient, the purified immune cells were collected from the interphase in a volume of 1.5 mL. After washing in 13 mL PBS/FCS and centrifugation at 800*×* g for 8 mins, cells were resuspended in an appropriate volume and transferred to the staining plate to perform flow cytometry staining.

#### Isolation of other organ immune cells

**Spleens** were brayed with the plunger of a syringe in 1 mL PBS/FCS and filtered through a 100 µm mesh into a 15 mL tube. 10 mL red blood cell lysis buffer was added. After 8 min of incubation, cells were centrifuged for 8 min at 400*×* g. The supernatant was discarded, and cells were washed once in 5 mL PBS/FCS and centrifuged for 5 min at 400*×* g before flow cytometry staining. **Celiac, portal and inguinal lymph nodes** were dissected from the tissue, and remaining fat was removed. Peyer’s patches were removed from the small intestine by cutting them out with ine scissors and collected in PBS/FCS. Lymph nodes and Peyer’s patches were then transferred and digested in 1 mL of RPMI containing 50 µg/mL Liberase*^TM^*(Roche) and 10 µg/mL DNase (Roche) in 10 % FCS/RPMI for 40 min in a shaking water bath at 37 °C. After 20 min of incubation, the tissue was mechanically disrupted by pipetting up and down. At the end of the incubation, cells were filtered through a 100 µm filter (Falcon), washed, and then centrifuged for 5 min at 400*×* g before flow cytometry staining. **Blood** was collected in EDTA-coated tubes (Sarstedt) filled with 500 µL of 20 mM EDTA in PBS. For red blood cell lysis, the cell suspension was transferred into 10 mL RBC lysis buffer (150 mM NH_4_Cl, 10 mM KHCO_3_, 0.1 mM EDTA-Na_2_ in distilled water [dH2O]), mixed and incubated for 8 min at room temperature (RT). Cells were immediately centrifuged for 8 min at 400*×* g, washed in 5 mL PBS/FCS, and centrifuged again for another five minutes at 400*×g* before staining. **Bone marrow** cells were isolated by dissection, cleaning and flushing it out from one tibia for adult mice or both tibiae for neonatal mice using 2 mL of PBS/3 % FCS. The cell suspension was centrifuged for 5 min at 400*×*g, then resuspended in 2 mL red blood cell lysis buffer and incubated for 2 min at room temperature to lyse erythrocytes. Next, cells were centrifuged (5 min at 400*×*g, 4C), washed twice in PBS/FCS and stained. **Small intestinal lamina propria:** After removal of the small intestine from the sacrificed mouse the adhering fat was removed and the intestine was cut open longitudinally. Intestinal content was removed by shaking in 10 mL PBS in a petri dish. The intestinal tissue was then shaken at 37 °C and 225 rpm for 20 min in 2 mM EDTA in 20 mL Hank’s balanced salt solution (Gibco)/3 % FCS, shaken vigorously by hand at the end of the 20 min incubation period and then filtered through a 100 µM cell strainer (Corning). The intestinal tissue remaining in the filter was transferred to a new tube with 2mM EDTA in 20 mL Hank’s balanced salt solution (Gibco)/3 %FCS and the incubation step was repeated. The remaining intestinal tissue was cut into small pieces and then digested for 40 min at 37 °C and 225 rpm in a shaking water bath in 10 mL RPMI/10 %FCS containing LiberaseTM (Roche; 30 µg enzyme/ mL digestion solution) and DNase (Roche) (10 µg enzyme/ mL digestion solution). 20 min into the incubation period, tubes were additionally shaken by hand to improve tissue disintegration. After 40 min, the digestion was stopped by adding 30 mL of PBS/3 %FCS and cells were centrifuged for 10 min at 800 x g and the supernatant was discarded. For enrichment of immune cells, a density gradient was performed similar to the liver immune cell isolation. **Colon lamina propria:** After removal of the colon from the sacrificed mouse, the adhering fat and faeces were removed, and the colon was cut open longitudinally and washed in a petri dish with 15 mL cold HBSS/3 % FCS. The colon tissue was then shaken at 37 °C and 225 rpm for 20 min in 2 mM EDTA in 20 mL HBSS/3 % FCS, shaken vigorously by hand at the end of the 20 min incubation period, and then filtered through a 100 µM cell strainer to ensure the removal of epithelial cell lining. This was repeated twice in total. The remaining colon tissue was cut into small pieces and then digested in Collagenase D (12.5 µg/ml, Roche), Dispase (10µg/ml, Gibco)) and DNase (10 µg/mL; Roche) in 10 mL RPMI/3 % FCS at 37 °C for 45 min. At the end of the incubation period, tubes were vigorously shaken by hand to improve tissue disintegration, filtered through a 100 µM cell strainer, and then directly centrifuged for 8 min at 400*×*g at 4 °C. For enrichment of immune cells, a density gradient was performed similar to liver immune cell isolation.

### Flow cytometry staining

After obtaining a single cell suspension, cells were transferred into a 96 well plate for staining and incubated with TruStainFcx (1/100 diluted PBS/FCS; Biolegend) for 10 min at 4 °C to block unspecific binding by the macrophage receptors CD16 and CD32. For CCR7 detection on T cells, CCR7 antibody was added 1:200 and cells were incubated 1 h at 37 °C. Cells were centrifuged at 400 x g, 4 °C for 5 min. Fixable viability dye 780 (ebioscience; 1/1000 diluted in PBS), Zombie UV or ZombieNIR fixable viability dye (Biolegend; 1/1000 diluted in PBS) stained for 30 min at 4 °C in a total volume of 100 µL per well, then cells were centrifuged at 400 x g, 4 °C for 5 min. Surface staining was done in 50 µL of antibody staining mix for 20-30 min at 4 °C. Cells were fixed with Foxp3 staining kit (ebioscience; for detection of transcription factors) or with Cytofix/Cytoperm kit (BD Biosciences; for the concomitant detection of cytosolic proteins such as cytokines or GFP with transcription factors) for 20 min at RT according to the manufacturer’s instructions. Cells were washed twice in the respective Perm Buffer. Intracellular staining was done in the Permeabilization Buffer of the respective kit overnight (14-18 h at 4 °C). Cells were washed once in PBS/FCS and then resuspended in PBS/FCS containing 5 % Precision Count Beads (Biolegend) for determination of cell number measured by the flow cytometer. Samples were acquired on a 5-laser spectral cytometer (Cytek Aurora, Cytek Biosciences) using spectral unmixing with autofluorescence extraction. Data were unmixed with SpectroFlo (Version 3.3) and analysed using FlowJo (Version 10, LLC).

Antibodies used for **surface staining and T cell phenotyping** were CD8α-SparkUV387 (Clone: 53-6.7, BioLegend), CD8α-BUV395 (Clone: 53-6.7, BD Biosciences), Bst2-BUV395 (Clone: Y129, BD Biosciences), TCRβ-BUV496 (Clone: H57-597, BD Biosciences), NK1.1-BUV563 (Clone: PK136, BD Biosciences), KLRG1-BUV661 (Clone: 2F1, BD Biosciences), CD4-BUV737 (Clone: RM4-5, BD Biosciences), CD45.1-BUV737 (Clone: A20, BD Biosciences), CD11c-BUV737 (Clone: N418, BD Biosciences), CD11c-BUV805 (Clone: N418, BD Biosciences), hCD2-BV421 (Clone: RPA-2.10, BioLegend), hCD2-PE (Clone: RPA-2.10, BioLegend), XCR1-BV421 (Clone: ZET, BioLegend), XCR1-BUV805 (Clone: ZET, BD Biosciences), TCR Vα2-BV421 (Clone: B20.1, BioLegend), PD-1-BV510 (Clone: 29F.1A12, BioLegend), CD90.2-BV570 (Clone: 30-H12, BioLegend), CD73-BV605 (Clone: TY/11.8, BioLegend), TCRδ-BV605 (Clone: GL3, BioLegend), CD69-BV650 (Clone: H1.2F3, BioLegend), CD69-BV785 (Clone: H1.2F3, BioLegend), Vsig4-BV605 (Clone: JAV4, BD Biosciences), CD44-BV711 (Clone: IM7, BioLegend), CD44-BV785 (Clone: IM7, BioLegend), TCR Vα2-FITC (Clone: B20.1, BioLegend), TCRδ-FITC (Clone: GL3, BioLegend), CD209α-FITC (Clone: MMD3, BioLegend), MHCII-SparkBlue550 (Clone: M5/114.15.2, BioLegend), MHCII-PE-Cy7 (Clone: M5/114.15.2, BioLegend), Nrp-1-RealBlue780 (Clone: V46-1954, BD Biosciences), CCR7-PE (Clone: 4B12, BioLegend), CCR7-PE-Cy5 (Clone: 4B12, BioLegend), CXCR6-PEDazzle594 (Clone: SA051D1, BioLegend), CD64-PEDazzle594 (Clone: W18349C, BioLegend), CD62L-PE-Cy5 (Clone: MEL-14, BioLegend), F4/80-PE-Cy5 (Clone: BM8, BioLegend), FR4-PE-Cy7 (Clone: 12A5, BioLegend), PD-1-PE-Cy7 (Clone: 29F.1A12, BioLegend), CXCR3-APC (Clone: CXCR3-173, BioLegend), CD45-R718 (Clone: 30-F11, BD Biosciences), CD45.2-AlexaFluor700 (Clone: 104, BioLegend), and CD4-APCFire810 (Clone: GK1.5, BioLegend).

Antibodies used for **nuclear staining** were Foxp3-eFluor450 (Clone: FJK-16s, Invitrogen), Foxp3-PE (Clone: FJK-16s, Invitrogen), Rorγt-BV480 (Clone: Q31-378, BioLegend), Ki-67-RealBlue705 (Clone: B56, BioLegend), Ki-67-FITC (Clone: B56, BioLegend), Ki-67-PE-efluor610 (Clone: SolA15, Invitrogen), and Helios-PE (Clone: 22F6, BioLegend). Antibodies used for **cytosolic staining** were IFNγ-PE (Clone: XMG1.2, BioLegend) and IL-17-AlexaFluor647 (Clone: TC11-18H10.1, BioLegend).

### Sequential multiplex immunohistochemistry

Sequential multiplex immunohistochemistry (mIHC) was performed as previously described(*53, 54*) The antibody elution buffer was prepared by mixing 675 μL distilled water, 125 μL 0.5 M Tris-HCl pH 6.8, 200 μL 10 % (w/v) sodium dodecyl sulfate, and 8 μL 2-mercaptoethanol. The following primary antibodies were used: anti-CD11c (clone D1V9Y, Cell Signaling, 1:500), anti-CD3e (clone E4T1B, Cell Signaling, 1:400), anti-CD4 (clone 4SM95, Thermo Fisher, 1:400), anti-CD8 (clone 4SM15, Thermo Fisher, 1:400), anti-Clec4f (clone 370901, R&D Systems, 1:1000), anti-Foxp3 (clone FJK-16s, Thermo Fisher, 1:200), anti-IBA1 (polyclonal, VWR #100369-764, 1:1000), anti-Lyve-1 (polyclonal, Abcam #ab14917, 1:200), anti-MHC-II (clone M5/114.15.2, BioLegend, 1:200), anti-PD-1 (polyclonal, R&D Systems #AF1021, 1:200), and anti-PD-L1 (clone E1L3N, Cell Signaling, 1:200).

Secondary antibodies included Alexa Fluor® 647–conjugated anti-rabbit IgG (H+L), F(ab’)₂ fragment (Cell Signaling #4414, 1:1000), Alexa Fluor® 750–conjugated goat anti-rat IgG H&L (Abcam #ab175751, 1:500), Alexa Fluor™ 750–conjugated goat anti-rabbit IgG (H+L), cross-adsorbed (Invitrogen #A-21039, 1:500), and Alexa Fluor® 647–conjugated anti-rat IgG (H+L) (Cell Signaling #4418, 1:1000).

Image analysis was performed with an in-house optimized image processing pipeline and software tools.(*53*) Clusters were defined as consisting of at least 3 cells of each cell type (CD3+ or CD11c+IBA1-Clec4f-) clustering in direct proximity and counted manually across the whole slides.

### Immunofluorescence imaging of CCR7KO and WT mice (Ext, Data 7a)

Samples were fixed in 4 % PFA, embedded and cryosectioned. Samples were blocked with serum and stained with aCD3 (500A2, hamster) – FITC 1:200 and 1:1000 DAPI overnight, followed by a wash step in >5ml wash/stain buffer for >4h at 4°C and constant agitation and mounted on glass slides with ProLongGold.After imaging, CD3+ cells were counted manually and in a blinded manner by 3 different persons. The mean of the results of the three counting persons was plotted for each slide.

### scRNASeq experiment and analysis

Liver cells were isolated as detailed in the isolation section. All pipetting steps were performed with wide-orifice low-binding pipets (VWR), After collecting the pellet from the Tissue Grinder device (FFX Technologies), cells were resuspended in 50 mL PBS/FCS and hepatocytes were pelleted by centrifugation at 50 x g for 2 min at 4 °C. The supernatant was centrifuged for 8 min at 400 x g at 4 °C to pellet the fraction enriched for immune cells. The pellet was resuspended in PBS/1 % BSA, cells were counted and stained for 30 min with the corresponding TotalSeqC-Hashtag antibody 1-3 (Biolegend) at 4 °C at a dilution of 1:3200 in 50 µL PBS/1 % BSA per million cells.After incubation, 5 mL of PBS/0.04 % BSA were added to the cell suspension and cells were pelleted by centrifugation for 5 min at 400 x g. The pellet was resuspended in 5 mL of PBS, incubated for 5 min at RT, then transferred to a new tube, centrifuged down and washed twice PBS/1 %BSA. Cells from the three replicates were then pooled and counted again. TotalSeqC-antibodies (Biolegend) for protein detection (TotalSeq-C1058, C0077, C0073, C0004; BioLegend) and flow cytometry staining antibodies (CD4-PECy7, Clone: GK1.5, BioLegend, TCRb-APC, Clone: H57-597, Biolegend, CD45-FITC, Clone: 30-F11, Biolegend) were added in their respective dilutions and stained for 30 min at 4 °C. After incubation, 5 mL of PBS/0.04 % BSA were added to the cell suspension and cells were pelleted by centrifugation for 5 min at 400 x g. The pellet was resuspended in 5 mL of PBS, incubated for 5 min at RT, then transferred to a new tube, centrifuged down and washed once with PBS/1 %BSA. DAPI was added for detection of dead cells and incubated for 2 min at 4 °C. Then, cells were centrifuged down for 5 min at 400 x g and resuspended in PBS/FCS for FACS sorting. Cells were sorted at the Aria II or Aria Fusion sorter (BD Biosciences) into 1.5 mL tubes with RPMI/10 % FCS at 4°C to collect at least 20.000 CD4 T cells and 200.000 other immune cells. Cells were then washed with PBS/FCS, counted again and adjusted to a concentration of 1.000 cells/µL. CD4 T cells and other immune cells were pooled in ratio of 1 + 3 aiming for a cell recovery of 2.500 CD4 T cells and 7.500 other immune cells per sample. Then, cells were put on the Next GEM Chip K to initiate the library generation workflow (10X Genomics). Single cell RNA sequencing workflow was performed with the 5’ Immune Profiling with Feature Barcoding Kit (10X Genomics) according to the manufacturer’s instructionsLibraries were quality controlled at the TapeStation (Agilent) sequenced on the Illumina NovaSeq system with an SP (for shallow sequencing to determine library composition) or S1 cartridge and preprocessed via the Cell Ranger pipeline by the Genomic Facility at IZKF Aachen.

Raw FASTQ data was analysed with the “CellRanger” pipeline (10X Genomics) and then in the R package Seurat(*55*) (v5.0.3.) to process all data: To demultiplex the biological replicates by hashtag antibodies, generate a separate data assay for hashtag antibodies along with the transcriptomics assay, we applied the function “HTODemux”. Benchmarking several options for “quantile” yielded the cutoff 0.99 for an optimal separation of cells per mouse. Cells assigned to more than 1 hashtag were classified as doublets and removed from the dataset. Cells predicted to be negative for hashtags were only removed for analyses of individual mice, e.g. in clonotype analysis, but left in the data for reclustering and DE gene analysis. Ambient RNA contamination was predicted by “decontX” from the “celda” package(*56*) (v1.18.2). Few remaining potential doublets were predicted and removed by the package DoubletFinder(*57*)(v2.0.4). For quality control, cells containing more than 5 % mitochondrial RNA counts, with a complexity (log10(nFeature_RNA) / log10(nCount_RNA)) < 0.8 or an ncount of > 2.5 * IQR above the 75^th^ quantile of the whole sample were removed.

Cell cycle scores for all cells were obtained by the built-in function “CellCycleScore” in Seurat and regressed out, as well as TCR genes (Trav and Traj genes), meaning they were not used for any clustering processes, while remaining in the data for DE gene comparisons. PCA and UMAP dimensional reduction were applied and clustering was performed using the built-in function RunPCA, RunUMAP, FindNeighbors and FindCluster in Seurat with a cluster resolution of 0.5. Data from different samples were integrated using the CCA method built-in Seurat. DE genes between clusters were found by conducting a Wilcoxon test with Benjamin-Hochberg correction. Ligand receptor analysis was performed by CellChatDB(v2.1.2, PMID: 39289562). Fig.s were prepared using Seurat and ggplot2 (v3.5.2).

To recluster only CD4 T cells, first all clusters containing T cells/ILCs and NK cells from Fig. S3c were reclustered (Fig. S3d) and then only CD4+ T cell containing clusters were reclustered to obtain the CD4 T cell UMAP (Fig. S3e) and merged into clusters comparable to flow analysis, using the following gene modules: naïve (Ccr7, Sell, Tcf7), Treg (Foxp3, Ikzf2, Ctla4), activated Tconv (Cxcr3, Id2, Cd44, Mki67) and NKT (Klrb1c)) These clusters were then used for DE gene analysis. Heatmaps were generated by normalizing gene expression per gene from the Seurat object for each mouse.

For VDJ analysis, individual mice were separated by hashtag and the package SC repertoire was used. R20 and R50 values were calculated according to Connors et al (add ref). To recluster only DC, only myeloid cell clusters were chosen from the UMAP in Fig. S7a, reclustered, manually annotated according to cluster marker and canonical genes and only DC containing clusters were chosen. Clusters were merged and a T cell contamination was removed (Fig. S6f), resulting in the UMAP shown in Fig. 3c.

Raw FASTQ data has been deposited in ENA with accession code PRJEB104713 and under the Zenodo ID (DOI:10.5281/17940609

### Xenium spatial transcriptomics

For 10X Xenium targeted spatial transcriptomics, we designed a 480-plex custom panel based on our scRNASeq data and the Liver Cell Atlas^56^. Sample processing was done on 5 µm thick paraffine embedded liver sections as described by the manufacturer using the Cell Segmentation kit and Xenium V1 chemistry. Analysis of the data was conducted using the SPARROW pipeline(*58*). The main steps consisted of image pre-processing, nuclei segmentation, transcript registration based on the new segmentation, cell type annotation, and cell neighbourhood analysis.

Image pre-processing was performed on the DAPI channel with the aim to improve the segmentation results. We applied a min-max filter with a size of 35 pixels to subtract background noise, and contrast enhancement using the CLAHE algorithm with a clip limit of 3.5. Nuclei segmentation was then performed on the pre-processed image using the cellpose model(*59*), which is a U-Net-like convolutional neural network (depth of 50 pixels and reflective boundary for the mapping overlap). The cellpose model was applied using an estimated nucleus diameter of 7 pixels; flow error threshold of <0.65, minimum area of 50 total pixels for segmented objects; probability threshold of >0.0; size fraction <0.4 0.4. The parameters for both the image pre-processing and segmentation were chosen based on grid searches and visual examination of the results. A combination of 25 different parameters for pre-processing and 156 for the segmentation were scanned.

Transcript registration was done using the new nuclei segmentation layer and the transcriptomics data was normalised based on the nuclei sizes. We removed the *Isg15*, *Top2a, Birc5, Ube2c, Cenpa* and *Mki67* genes for the downstream analysis, as these genes were strongly driving the clustering.

Leiden clustering was performed using the default parameters(*60*). We looked at the top 20 genes of each cluster and annotated DCs and T cells the clusters base on expression of canonical marker genes. Extended Data 7b displays the results of the nuclei segmentation highlighting the DCs and T cells that were identified using Leiden clustering. In order to formally define DC-T cell clusters, we applied the DBScan algorithm implemented by the scikit-learn Python library (v1.7.1) using the centroid coordinates of the annotated DCs and T cells (Fig. S7c). DE gene analysis was conducted for DCs and T cells in and outside clusters (Figs. S7f&i): Volcano plots display the results marking the top 10 DE genes. As close proximity of the cells leads to detection of transcripts from neighbouring cells (e.g. MHCII genes in T cells and CD3 in DCs) even with our very conservative nuclei segmentation, we filtered the DE labelled genes for T cell/DC genes from our panel only for Fig. 3j.

The raw and processed 10X Xenium data are available in Zenodo, ID17940609:

### AI-assisted manuscript preparation

ChatGPT (OpenAI; model: GPT-4.5/5.1 Thinking) was used during manuscript preparation to assist with language editing (clarity, concision, and organization) and to generate alternative phrasing. The tool was not used to generate or analyze primary data, to perform statistical analyses, or to generate references/citations. All AI-assisted text was reviewed, edited, and verified by the authors.

**Supplementary figure 1:**
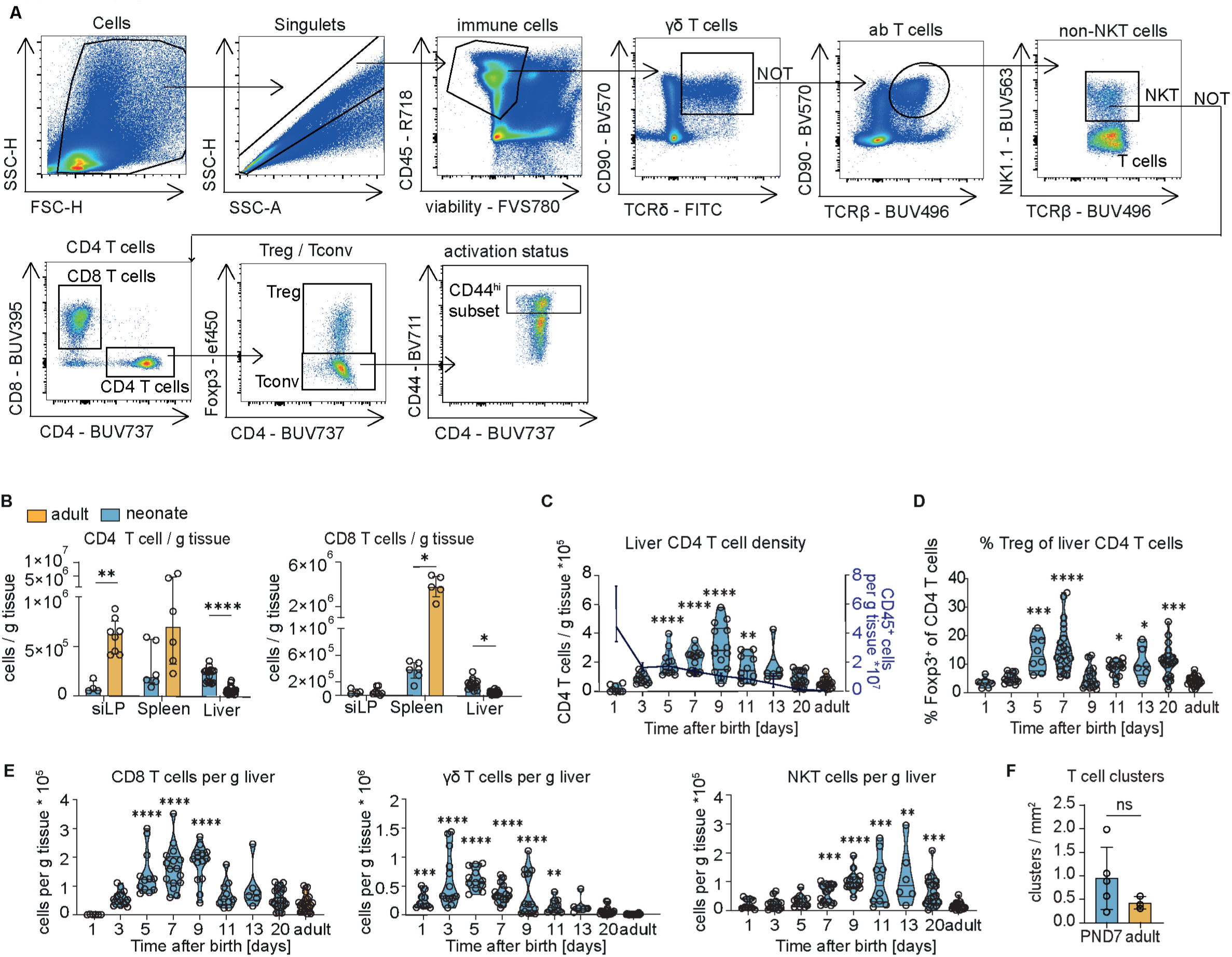
Flow cytometric quantification of T cells in the neonatal liver. (A) Gating strategy for the identification of T cell subsets and phenotype used in all flow cytometry experiments (if not indicated otherwise). (B) Cell densities of CD4 and CD8 T cells in liver tissue in small intestinal lamina propria (siLP), Spleen and liver. Multiple Mann Whitney U test. 4-1O mice from 1-2 independent experiments. (C) Kinetics of CD4 T cells (violin plots) and total CD45^+^cell density (line showing median + IQR) in the neonatal liver. Kruskal Wallis test + Dunn’s comparison of each lime point to adult group. Violin plots of 6-20 mice per timepoint from 2-6 independent experiments. (D) % of Tregs (Foxp3^+^) within CD4 T cells in the neonatal liver. Kruskal Wallis test + Dunn’s comparison of each lime point to adult group. Violin plots of 6-20 mice per limepoint from 2-6 independent experiments. (E) Kinetics of CD8 T, ydT and NKT cell density in the neonatal liver. Kruskal Wallis test + Dunn’s comparison of each lime point to adult group. Violin plots of 6-20 mice per timepoint from 2-6 independent experiments. (F) Ext. Data 1f: Density of manually counted microclusters of CD3^+^ cells per slide in neonatal (7-day-old) and adult WT livers (3-5 mice per group). Unpaired I-test. Asterisks indicate significance levels: * < 0.05, ** < 0.01, *** < 0.001 **** < 0.0001. All tests were performed in a two-sided manner.

**Supplementary figure 2:**
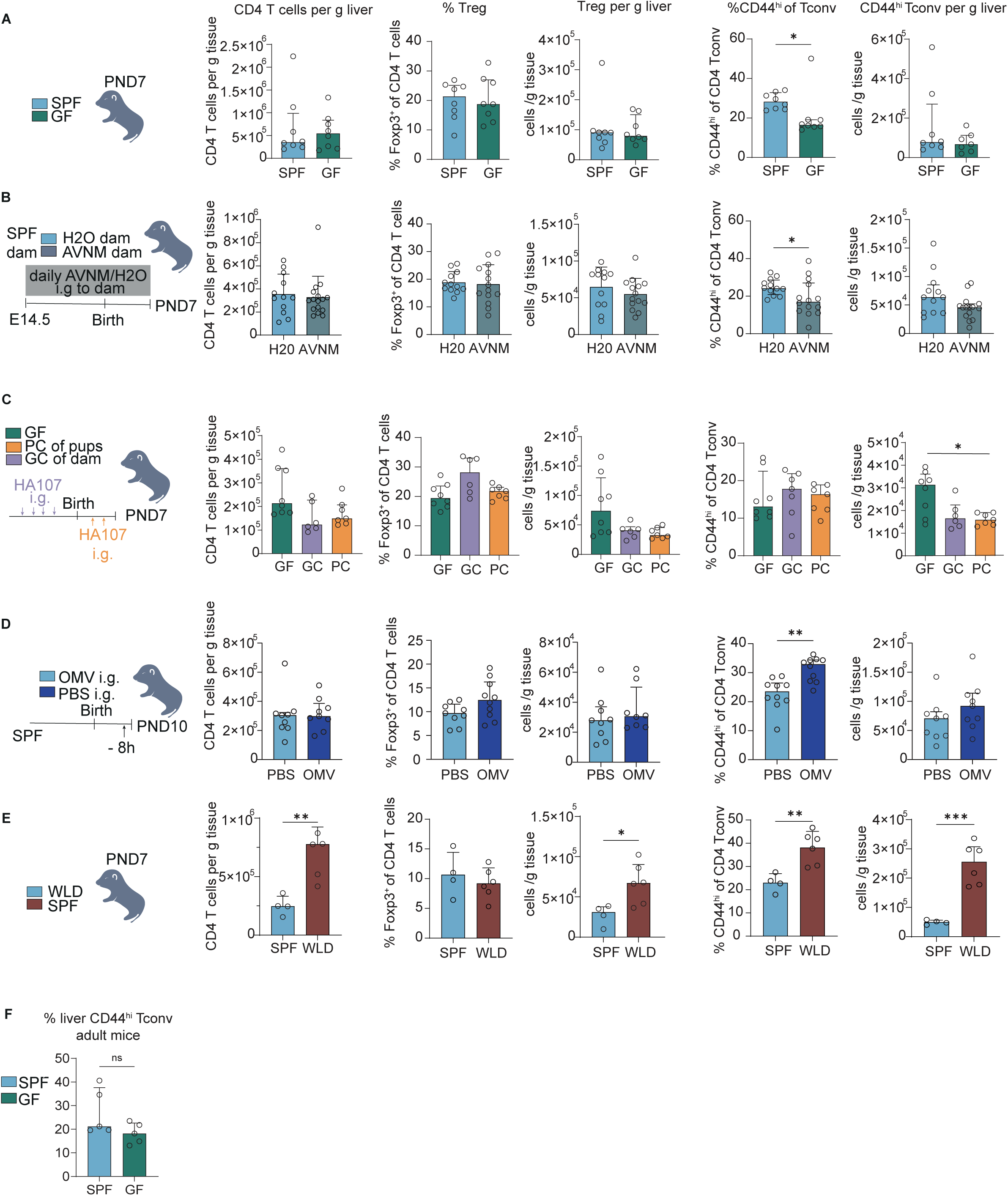
Experimental design and CD4T cell density, % Treg of CD4 T cells, Treg density, % CD44^hi^ Tconv and CD44^hi^ Tconv density in livers of neonatal mice in different microbial exposure models: (A) SPF vs germ-free (GF) mice: 8 mice pooled from two independent experiments per group. Mann Whitney U tests or unpaired t-test ( % Tregs of CD4 T cells). (B) Daily treatment of dams from E14.5 until PND7 with broad-spectrum antibiotics (Ampicillin, Vancomycin, Neomycin ad libitum in drinking water and daily administration of Metronidazol i.g.) 12-15 mice from 2 experiments. Mann Whitney U tests or unpaired t-test (Tregs per g liver) or Welch t-test ( % Tregs of CD4 T cells) (C) Gestational colonisation of germ-free dams (GC) and postnatal colonisation of germ-free pups (PC) by i.g. gavage with the auxotrophic E. coli strain HA107. 6-8 mice from 2 independent experiments (1 experiment for PC). Brown-Forsythe ANOVA + Holm-Sidaks’post hoc test ( % Treg, Treg density), One Way ANOVA + Holm-Sidak multiple comparison ( % CD44^hi^ Tconv) or Kruskal Wallis test + Dunn’s (CD4 T cell density and CD44^hi^ Tconv density). (D) Gavage of SPF pups at PND10 with E. coli Outer Membrane Vesicles (OMVs) or PBS i.g.. 8-10 mice from 2 independent experiments. Unpaired t-tests (all data sets except CD4 T cell density (Mann Whitney U test) (E) SPF vs wildlings: 4-6 per group mice from 1 experiment. Unpaired-tests (CD4 T cell density, Treg density, % Treg) or Welch t-tests (Cd44^hi^ % and density) (F) % CD44^hi^ of Tconvs in livers of 8-10 week old SPF and germ-free mice (GF). 5 mice per group from 2 independent experiments. Unpaired t-test. All experiments containing flow cytometry were done with the compared groups within 1 experiment processed on the same day. Asterisks indicate significance levels: * < 0.05, ** < 0.01, *** < 0.001 **** < 0.0001. All tests were performed in a two-sided manner.

**Supplementary figure 3:**
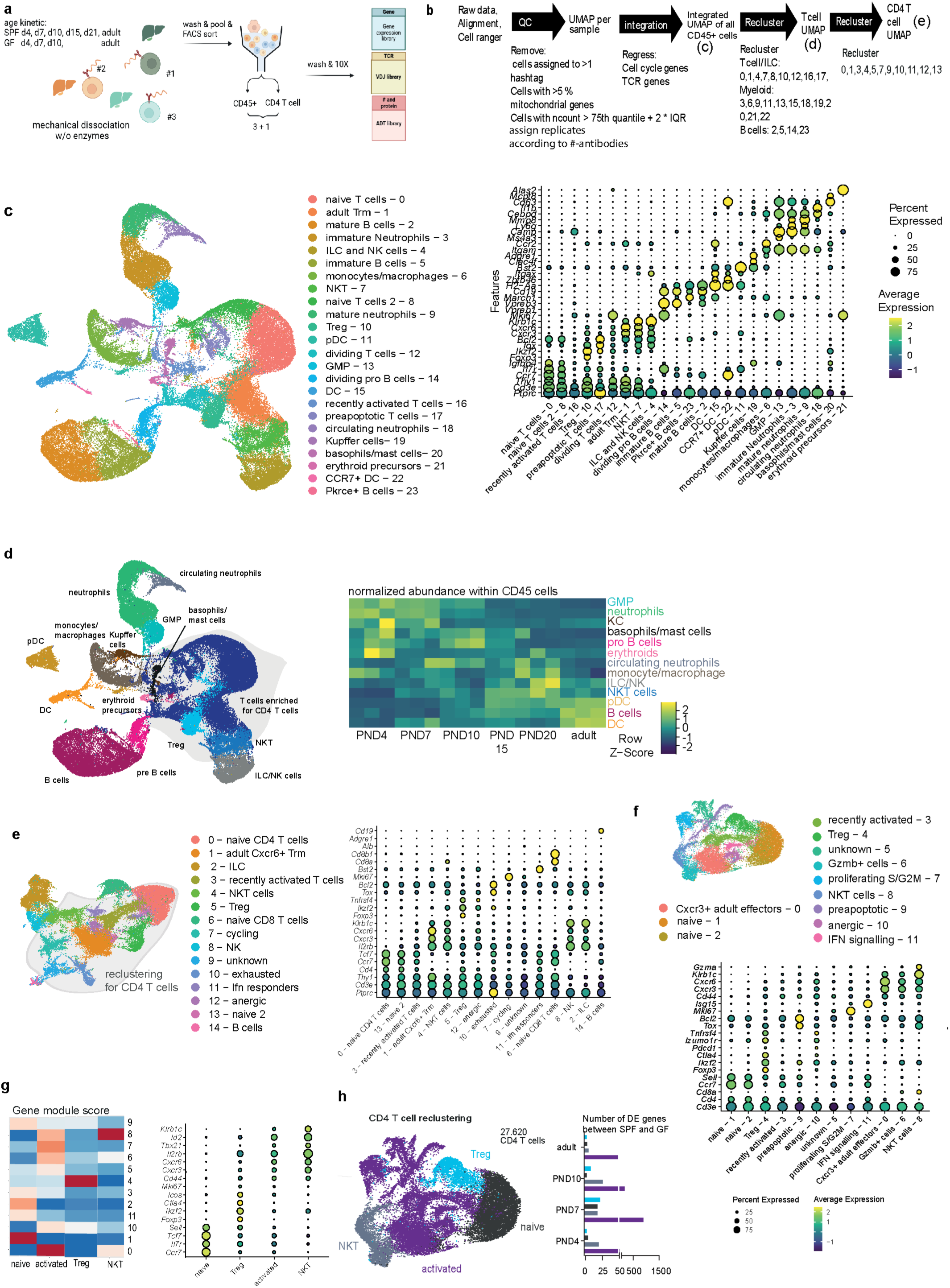
Analysis strategy of scRNASeq data and Extended Data. (A) Experimental design of scRNASeq experiment with TolalSeqC- hashtagged biological replicates (1 mouse per hashtag) and enrichment for CD4 T cells, Graphic created with Biorender, (B) Flow chart detailing QC metrics and reclustering strategy for scRNASeq data, (C) UMAP of all CD45+ cells, bubbleplot with canonical marker genes used for annotation; this UMAP was used for Cell Chat analysis in Figure 3, (D) UMAP of all CD45+ cells with merged clusters and heatmap of the z-score normalized % of total sample for each SPF mouse over the kinetic, (E) Reclustered UMAP of all T cells and ILCs and bubbleplot with canonical marker genes used for annotation, (F) Reclustered UMAP of CD4 T cells and bubbleplot with canonical marker genes used for annotation, (G) Gene modules and gene module scores of the clusters from (f) for merging of the clusters: naive: Ccr7, Sell, Tcf7; activated: CD44, Mki67, ld2; Treg: Foxp3, lkzf2, Ctla4; NKT: Klrb1c (H) Merged CD4 T cell UMAP and number of DE genes in each cluster between SPF and GF mice at different lime points after birth with bar plots color-coded by cluster,

**Supplementary figure 4:**
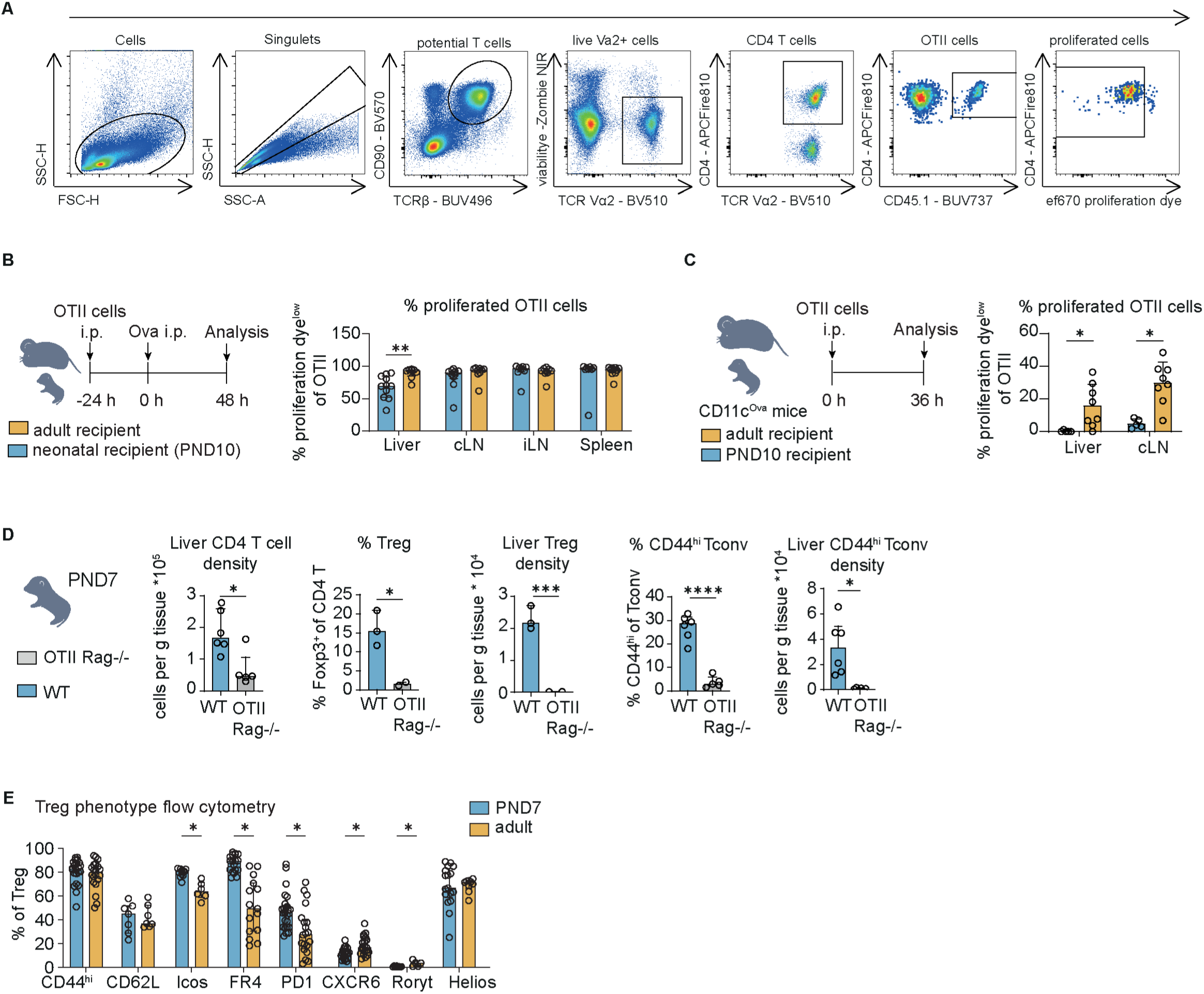
OTII experiments show delayed proliferation in neonatal liver tissue. (A) Gating strategy for identification of proliferated OTII cells for (b) and (c) (B) Experimental design of OTII transfer and intraperitoneal ovalbumin (ova) injection into neonatal and adult *WT* mice and % of proliferated cells of OTII cells in liver, celiac LN (cLN), inguinal LN (iLN) and spleen 48 h after antigen exposure. 8-1O mice from 2 independent experiments. Multiple Mann Whitney U test. + Holm-Sidak multiple test correction. (C) Experimental design of OTII transfer into neonatal and adult CD11c-Ova mice and % proliferated cells of OTII cells in liver and celiac LN (cLN) 48 h after antigen exposure. 7-9 mice from from 2 independent experiments. Multiple Mann Whitney U test + Holm-Sidak multiple test correction. (D) CD4 T cell density, Treg frequency and density, and activated Tconv frequency and density in the liver 7-day old B6 *WT* SPF or OTII RAG-/- mice (no exposure to the OTII antigen Ova in either of the groups. Barplots show median and IQR of 2 pooled experiments (1 experiment for Treg data) with 2-6 pups per group. Welch I-tests. (E) Treg phenotypic marker expression in PND7 and adult livers measured by flow cytometry. 6-19 mice per group from 3-5 independent experiments. Multiple Mann Whitney U test. All experiments containing flow cytometry were done with the compared groups within 1 experiment processed on the same day. Data show median and IQR if not otherwise indicated. Asterisks indicate significance levels: * < 0.05, ** < 0.01, *** < 0.001 **** < 0.0001 or not significant iflhere is no asterisk. All tests were performed in a two-sided manner.

**Supplementary figure 5:**
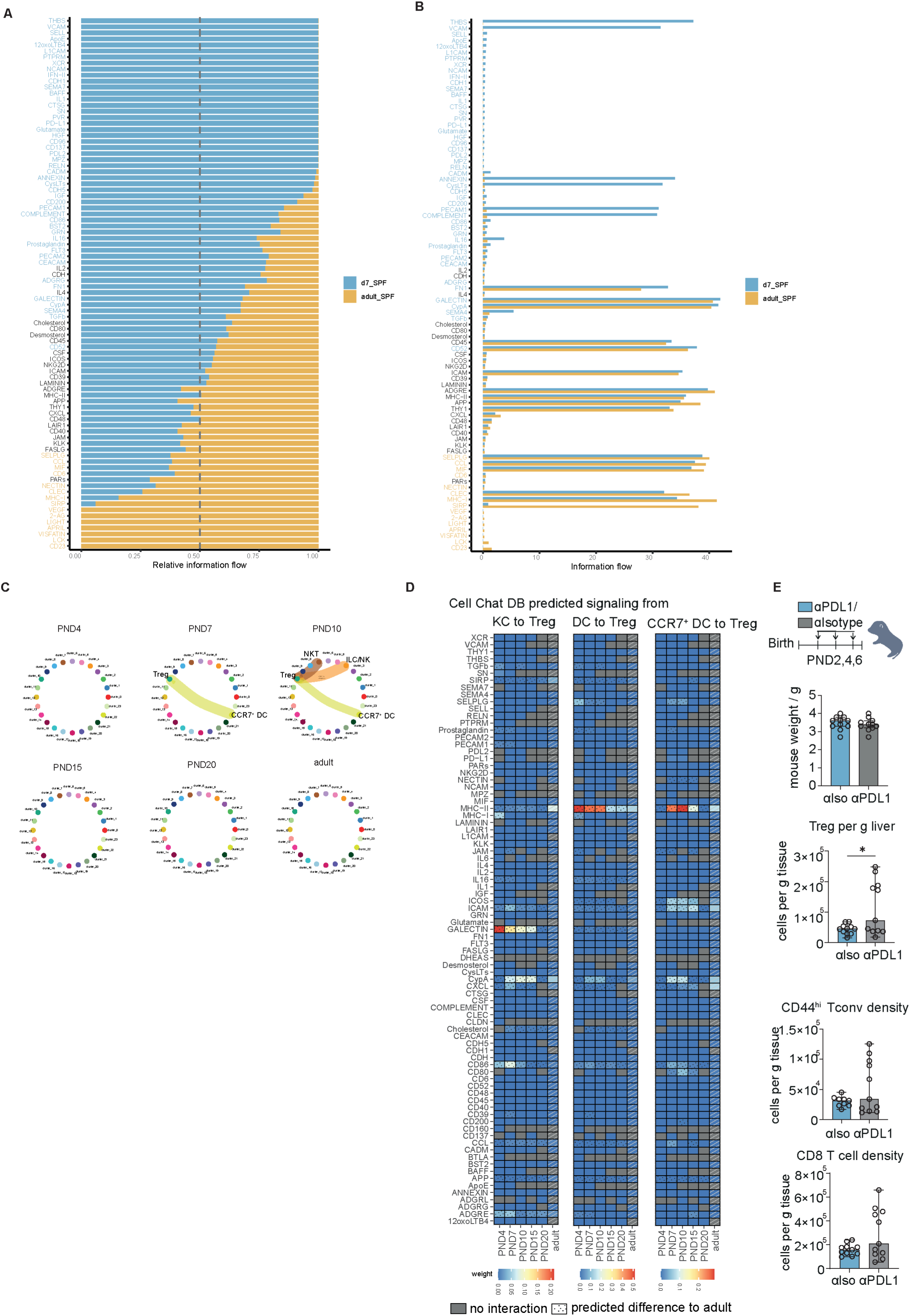
Cell Chat analysis and PDL1-blockade. (A) Comparison of relative information flow between neonatal (PND7) and adult liver total immune cells predicted by Cell Chat DB (blue = enriched signalling in neonate, yellow = enriched signalling in adult; black = no difference). (B) Absolute information flow between neonatal (PND7) and adult liver total immune cells predicted by Cell Chat DB. (blue = enriched signalling in neonate, yellow= enriched signalling in adult; black= no difference) (C) Circle plots indicating PD-L2 signaling in SPF at different limepoints after birth. Cluster indicate the clusters from Extended data 3c. (D) Heatmaps displaying all CellChat DB pathways signaling to the Treg cluster from either KC, DC or CCR7^+^ DC. (E) Experimental design for neonatal PD-L1 blockade, weights of aPD-L1 treated neonates and their littermates treated with alsotype antibody a PND7, Tregs per g liver, CD44^hi^ Tconvs per g liver and CD8 T cells per g liver. Unpaired I-test (mouse weight) and Mann Whitney U tests. 9-11 pups from 3 independent experiments. All experiments containing flow cytometry were done with the compared groups within 1 experiment processed on the same day. Data show median and IQR if not otherwise indicated. Asterisks indicate significance levels: * < 0.05, ** < 0.01, *** < 0.001 **** < 0.0001 or not significant if there is no asterisk. All tests were performed in a two-sided manner.

**Supplementary figure 6:**
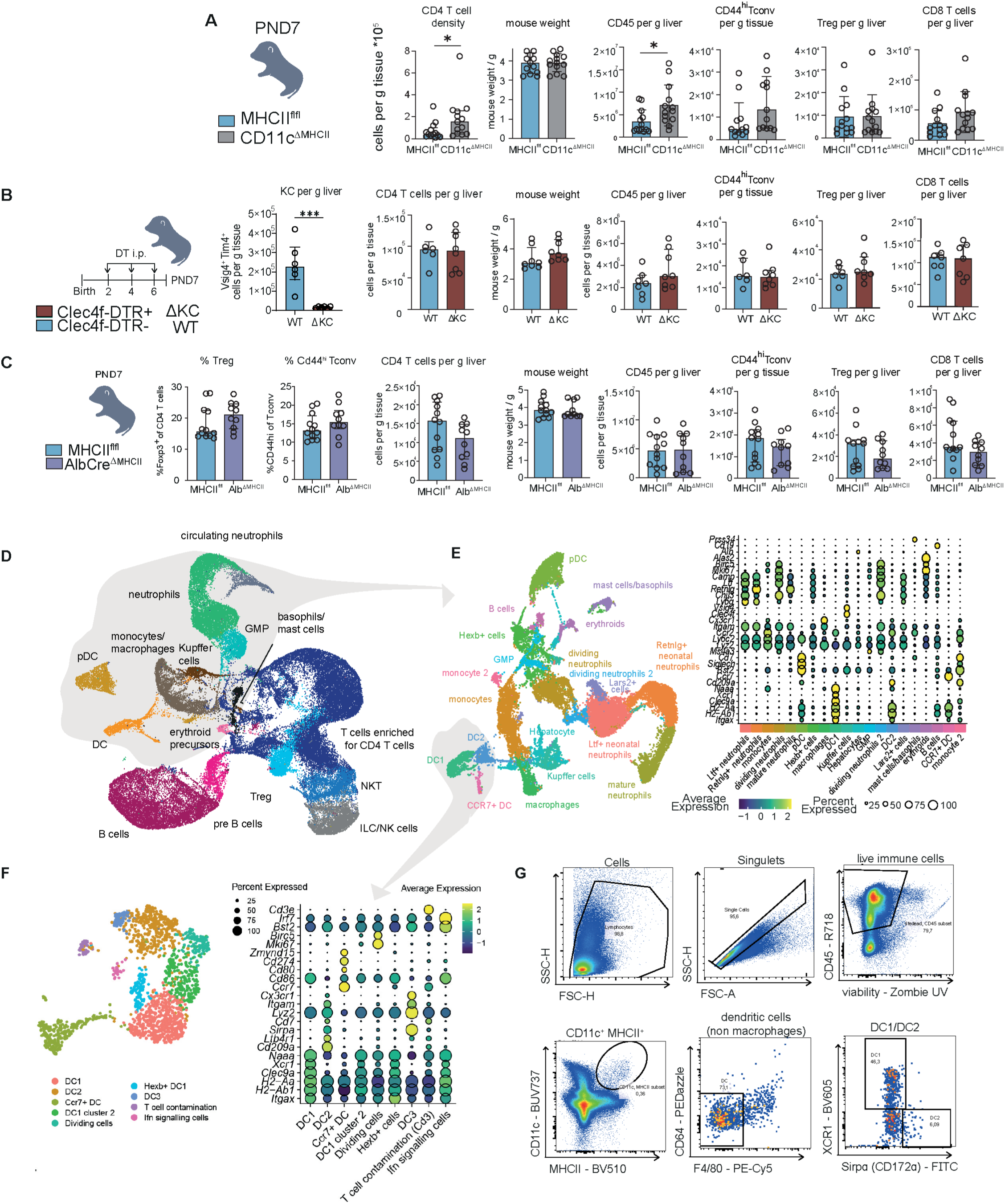
Flow cytometry of neonatal mice with conditional MHCII knock-out or APC depletion. (A) Extended flow cytometry data for neonatal CD11cCre MHCllflfl mice and their Cre-littermates. Liver CD4 T cell density, mouse weights at PND7, CD45+ cell per g liver, CD44^hi^ Tconvs per g liver, Tregs per g liver and CD8 T cells per g liver. 9-12 mice per group pooled from 4 independent experiments. Mann Whitney U tests. (B) Extended flow cytometry data for neonatal depletion of Kupffer cells. Clec4f-DTR + and Celec4f-DTR-littermates; KC density in liver tissue (Welch I-test); liver CD4 T cell density, mouse weights at PND7, CD45+ cell per g liver, CD44^hi^ Tconvs per g liver, Tregs per g liver and CD8 T cells per g liver (unpaired I-tests). 6-8 mice per group pooled from 2 independent experiments. (C) Liver CD4 T cell phenotypes in neonatal AlbCre MHCllflfl mice and their Cre negative littermates. 9-12 mice per group from 3 independent experiments. Mann Whitney U tests. Mann Whitney U tests. (D) CD45+ UMAP used for reclustering of all myeloid cells (gray circle). (E) Resulting Myeloid UMAP used for reclustering of DC (gray circle) and bubbleplot showing a selection of genes used for annotation. (F) DC UMAP used for analysis and bubble plot bubbleplot showing a selection of genes used for annotation. (G) Gating strategy for flow cytometry analysis of liver DC in neonatal and adult mice. All experiments containing flow cytometry were done with the compared groups within 1 experiment processed on the same day. Data show median and IQR if not otherwise indicated. Asterisks indicate significance levels:*< 0.05, ** < 0.01, *** < 0.001 **** < 0.0001 or not significant iflhere is no asterisk. All tests were performed in a two-sided manner.

**Supplementary figure 7:**
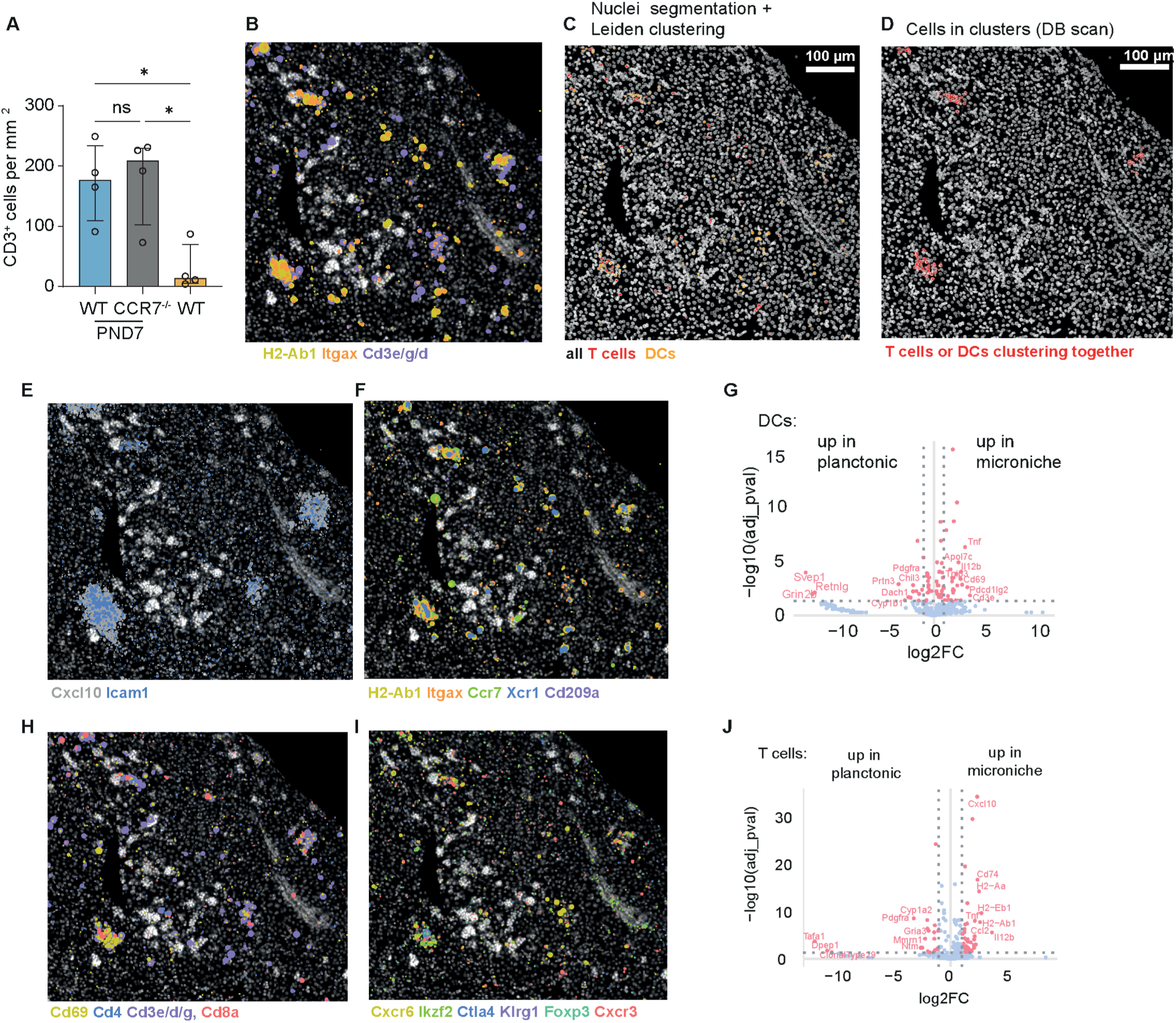
Ext. Data for Xenium spatial transcriptomics of neonatal liver. (A) CD3+ cells per mm’ in neonatal WT, CCR7 KO and adult WT mice, means of cell number manually counted in immunofluorescence microscopy pictures by 3 different persons in a blinded manner. Barplots show median + IQR of 4 mice per group. One Way ANOVA + Holm-Sidak’s multiple comparison. (B) Overview DAPI image + expression of DC *(ltgax, H2-Ab1)* and T cell transcripts (Cd3e, *CD3g, CD3d)* in 7-day old liver. (C) Segmented image with all DCs and T cells highlighted (D) Segmented image showing DCs and T cells within clusters defined using DBScan. (E) DAPI image + expression of *Cxcl10* and *lcam1* transcripts. (F) DAPI image + expression of DC *(ltgax, H2-Ab1)* and subsets (DC1: *Xcr1,* DC2: *Sirpa,* CCR7+ DC: *Ccr7)* (G) Volcano plot showing DE gene analysis of DCs inside and outside of microniches from Xenium panel. The top DE genes from the total panel are highlighted with labels. (H) DAPI image + expression of T cell marker transcripts *(Cd69* (activation), *Cd4, Cd3e, Cd3g, Cd3d, CdBa)* (I) DAPI image + expression of Treg *(Foxp3, lkzf2, Ctla4, Klrg1)* and tissue residency-associated transcripts *(Cxcr6, Cxcr3)* (J) Volcano plot showing DE gene analysis of T cells inside and outside of microniches from Xenium panel. The top DE genes from the total panel are highlighted with labels.

**Supplementary figure 8:**
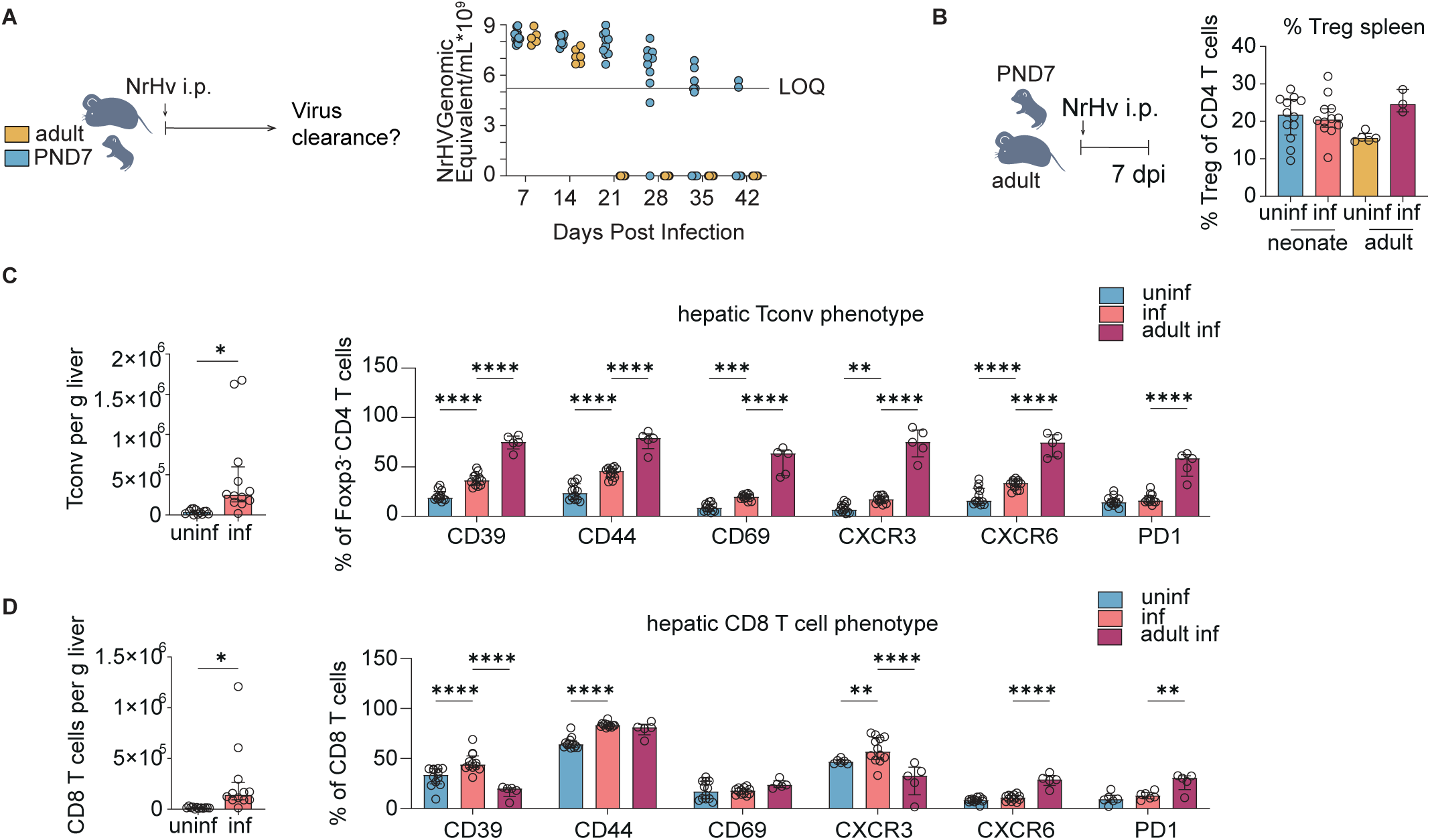
Extended data for analysis of NrHV infected mice. (A) Experimental design of neonatal NrHV infection. Viremia levels in neonatally or adult NrHV infected mice. 5-10 mice from 1 adult and 2 independent neonatal experiments. (B) Splenic Treg in NrHV-infected and uninfected neonatal and adult mice. Flow cytometry data of 11-12 neonatal mice from 2 pooled independent experiments and 3-5 adult mice from 1 experiment. One Way ANOVA + Tukey’s multiple comparison test. (C) Hepatic Tconv density and phenotypic markers in neonatal uninfected, neonatal infected and adult infected mice 7 days post-infection measured by flow cytometry. 12 neonatal mice per group from 2 independent experiments and 5 adult mice per group from 1 experiment. Two Way ANOVA with simple effects calculation between the neonatal infected group and the two other groups + multiple comparison correction via Holm-Sidak test. (D) Hepatic CDS T cell number and phenotypic markers in neonatal uninfected, neonatal infected and adult infected mice 7 days post-infection measured by flow cytometry. 12 neonatal mice per group from 2 independent experiments and 5 adult mice per group from 1 experiment. Two Way ANOVA with simple effects calculation between the neonatal infected group and the two other groups + multiple comparison correction via Holm-Sidak test.

**Supplementary figure 9:**
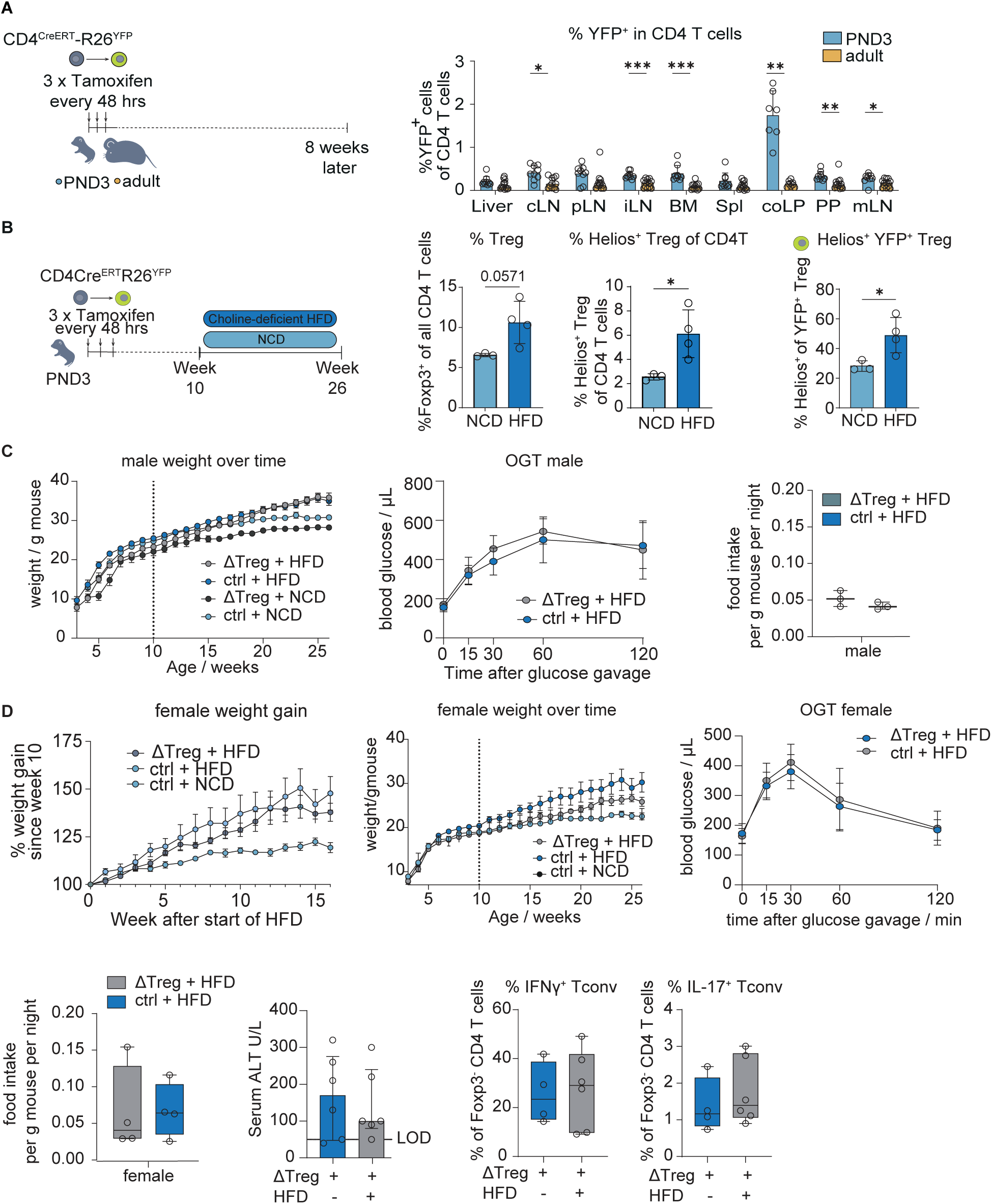
Timestamping and choline-deficient HFD experiments. (A) Experimental design and % YFP^+^ of CD4 T cells in different organs 8 weeks after timestamping. Multiple Mann Whitney U tests. (B) Experimental design of neonatal timestamping in combination with HFD. 1 representative of 2 experiments with littermates 3-4 mice per diet group. Unpaired t-tests. (C) Extended data for male mice on HFD after neonatal Treg depletion. Body weight over the course of the experiment, oral glucose tolerance test. Food intake per g body weight per night in the last week of HFD (3 mice per group pooled from 2 independent experiments). (D) Data for female mice on HFD after neonatal Treg depletion. Weight gain since start of HFD, body weight over the course of the experiment, oral glucose tolerance test. 7-8 mice per group from 6 independent experiments. Food intake per g body weight per night in the last week of HFD ((3 mice per group pooled from 2 independent experiments). 7 Serum ALT levels and flow cytometry measurement of IFNg and IL-17 producers in Tconvs after restimulation with PMA/Ionomycin. 4-6 per group from 4 independent experiments using only litters with both neoTreg and ctrl mice. Statistical analysis performed as was done for the male mice in the corresponding main figure.

